# Adhesion to nanofibers drives cell membrane remodeling through 1D wetting

**DOI:** 10.1101/393744

**Authors:** Arthur Charles-Orszag, Feng-Ching Tsai, Daria Bonazzi, Valeria Manriquez, Martin Sachse, Adeline Mallet, Audrey Salles, Keira Melican, Ralitza Staneva, Aurélie Bertin, Corinne Millien, Sylvie Goussard, Pierre Lafaye, Spencer Shorte, Matthieu Piel, Jacomine Krijnse-Locker, Françoise Brochard-Wyart, Patricia Bassereau, Guillaume Duménil

## Abstract

The shape of cellular membranes is highly regulated by a set of conserved mechanisms. These mechanisms can be manipulated by bacterial pathogens to infect cells. Human endothelial cell plasma membrane remodeling by the bacterium *Neisseria meningitidis* is thought to be essential during the blood phase of meningococcal infection, but the underlying mechanisms are unknown. Here we show that plasma membrane remodeling occurs independently of F-actin, along meningococcal type IV pili fibers, by a novel physical mechanism we term “ onedimensional” membrane wetting. We provide a theoretical model that gives the physical basis of 1D wetting and show that this mechanism occurs in model membranes interacting with model nanofibers, and in human cells interacting with model extracellular matrices. It is thus a new general principle driving the interaction of cells with their environment at the nanoscale that is diverted by meningococcus during infection.

## Main text

Control of the shape of biological membranes is fundamental for the maintenance of multiple functions in the eukaryotic cell^1^. Acting as the interface of the cell with its surrounding environment, the plasma membrane is a particularly important compartment that is subject to a precise control of its shape and dynamics. Plasma membrane remodeling occurs at very small scales, for example in the biogenesis of caveolae^2^ or during the formation of clathrin coated pits^3^. At larger scales, remodeling of the plasma membrane plays an important role in a wide variety of biological processes, such as the uptake of large particles by phagocytosis^4^ or in the formation of actin-based membrane structures that support cell migration and probing of the extracellular environment, such as filopodia or lamellipodia^5^. In the context of pathological conditions, especially in bacterial, viral and fungal infections, pathogens manipulate the shape of the plasma membrane to enter host cells. This is often achieved by diverting the actin cytoskeleton^6-8^. Other pathogens remain extracellular and must then resist mechanical strains such as those generated by flow^9^. The bacterium *Neisseria meningitidis* (or meningococcus) is a human pathogen that, while remaining extracellular^10^, massively remodels the host cell plasma membrane to form filopodia-like protrusions that intercalate between aggregated bacteria upon adhesion to the host cell surface. It was shown *in vitro* that plasma membrane remodeling allows *N. meningitidis* to proliferate on the outside of the host cell while mechanically resisting high shear stress levels^11^, suggesting a central role for plasma membrane remodeling in the blood phase of *N. meningitidis* pathogenesis where bacteria are subject to high shear. Colonization of the blood vessels by *N. meningitidis* eventually leads to a loss of vascular function that translates into hemorrhagic lesions in organs throughout the body, including the skin where it presents as characteristic purpuric rashes^12-14^. Despite the intensive use of antibiotics, the case fatality rate for meningococcal sepsis can still reach 52%^15^. Understanding this process is thus important in the study of both infectious processes and mechanisms of plasma membrane dynamics.

The molecular mechanisms by which *N. meningitidis* remodels the host cell plasma membrane are still elusive. While membrane protrusions are enriched in F-actin^16^, our previous work has shown that inhibition of actin polymerization^11, 16, 17^ or depletion of host cell ATP^17^ have no effect on the remodeling of the host cell plasma membrane. Bacterial type IV pili (T4P), which are long retractile fibers with a diameter of 6 nm, are required for plasma membrane remodeling in addition to their role in specific adhesion to human cells^12, 17^. Indeed, adhesion of non-piliated bacteria mediated by non-fibrillar adhesins, like Opa, does not lead to the formation of plasma membrane protrusions^19^. Furthermore, plasma membrane remodeling is tightly linked to the amount of T4P expressed by the bacteria, as a 30% decrease in T4P is sufficient to strongly decrease cell surface remodeling^20^. However, the molecular mode of action of T4P in plasma membrane remodeling is currently unknown.

Plasma membrane remodeling by meningococcus has only been observed in cultured cells. Therefore, we first sought to verify the existence of plasma membrane remodeling by meningococcus *in vivo*. Because of the human specificity of meningococcus, we used a recently developed humanized animal model of infection where human dermal vessels in skin is grafted onto immunodeficient mice^12^. Staining of human blood vessels during vascular colonization by *N. meningitidis* revealed that the surface of human endothelial cells was reorganized in the form of plasma membrane protrusions that intercalated between aggregated bacteria (Fig. 1a and Extended Data Fig. 1a and b), co-localized with T4P (Fig. 1a), and were enriched in F-actin (Extended Data Fig. 1a), paralleling earlier *in vitro* studies^11,16,17^. This data shows that the endothelial cell plasma membrane is remodeled during vascular colonization by meningococcus *in vivo*, supporting the idea of a key role of plasma membrane remodeling in meningococcal pathogenesis, and warranting to further study its underlying mechanisms.

**Figure 1.**
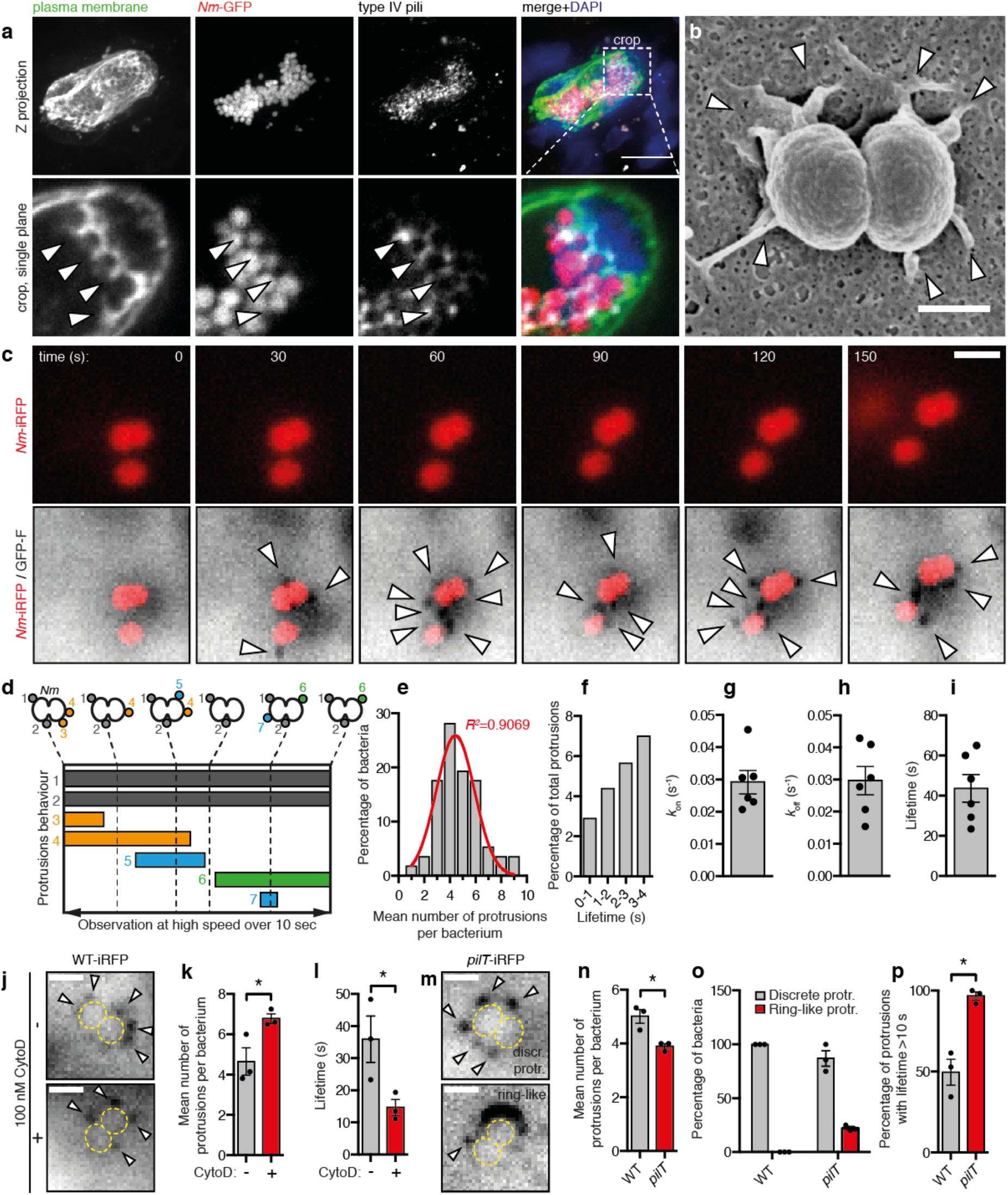
Plasma membrane remodeling by *Nm* occurs *in vivo*, and is initiated at the level of the individual bacterium *in vitro*, **(a)** Histoimmunolabeling of human blood vessels in a mouse after 3h of infection with *N. meningiiidis*(*Nm*)-GFP showing plasma membrane protrusions co-localizing with T4P between aggregated bacteria (arrowheads). Scale bar, **10** μm. Representative of **n=2** mice, **(b)** Scanning electron micrograph of an endothelial cell infected for 10 min. Arrowheads show plasma membrane protrusions. Scale bar, 500 nm. Representative of n=2 independent experiments performed in duplicate. **(c)** Oblique illumination live imaging of an endothelial cell expressing the membrane marker GFP-F (GFP-Farnesyl, inverted contrast) infected by individual Nm-iRFP. Arrowheads show dynamic plasma membrane protrusions. Scale bar, 2 μm. n>10 independent experiments. **(d)** Types of plasma membrane protrusions observed in high speed oblique illumination imaging. **(e)** Frequency distribution (gray bars) and Gaussian fit (red line) of the number of protrusions per bacterium. **(f)** Frequency distribution of the number of plasma membrane protrusions with short observed lifetimes. **(g-i)** Plasma membrane protrusions on-rate (*k*_on_), off-rate (*k*_off_) and mean lifetime. Gray bars and error bars represent the mean ±SEM over 57 bacteria in six independent experiments. Dots represent the mean values of 4-14 bacteria in each experiment. **(j-l)** Plasma membrane protrusions are still induced *Nm* in cytochalasin D-treated cells but have shorter lifetimes. Scale bars, 2 μm. Bars and error bars in graphs represent the mean ±SEM in three independent experiments. Dots represent the mean of each individual experiment. The total number of bacteria analyzed was 26 and 15 in non-treated and treated cells, respectively. **(m-p)** Mutant bacteria deficient for pilus retraction trigger plasma membrane protrusions with altered morphologies and that are no longer dynamic. Scale bars, 2 μm. Bars and error bars in graphs represent the mean ±SEM of three independent experiments. Dots represent the mean of each individual experiment. The total number of bacteria analyzed was 31 WT-iRFP and 28 *pilT*-iRFP.

At an early stage of infection, meningococcus adheres to the endothelium as individual bacteria^12^. Scanning electron microscopy (SEM) showed plasma membrane remodeling occurred even at the level of single bacteria as several discrete protrusions surrounding the bacterial body (Fig. 1b). It should be noted that in standard SEM technical conditions used here type IV pili filaments are not conserved. Live cell oblique fluorescence imaging of endothelial cells expressing a fluorescent plasma membrane marker revealed that these discrete protrusions were elicited immediately upon adhesion of individual bacteria to host endothelial cells and, while remaining in close proximity with the bacterial body, had dynamic properties (Fig. 1c and Movie S1). F-actin was not detectable in these protrusions over time ranges of 5-10 minutes, demonstrated by the absence of LifeAct-mCherry signal (Movie S2). Over several bacterial division events, plasma membrane protrusions accumulated in the nascent microcolony as bacteria proliferate on top of the host cell (Extended Data Fig. 1c and Movie S3). By recording images at high speed, we could track the fate of each plasma membrane protrusion (Fig. 1d-f). Individual bacteria induced 4 to 5 protrusions on average (Fig. 1e), some with lifetimes as short as 170 milliseconds (Fig. 1f). Within large bacterial aggregates, protrusions were no longer dynamic, and the rare disappearing events were restricted to the edge of the aggregate (Extended Data Fig. 1d and Movie S4), pointing to a possible maturation process stabilizing the interactions with the host cell. In individual bacteria, the growth and retraction rates of plasma membrane protrusions (*k*on and *k*_off_, respectively) were similar, indicating that there was a dynamic equilibrium between appearance and disappearance of the protrusions (Fig. 1g and h). The average lifetime for plasma membrane protrusions, equal to 1/*k*_off_, was 44 ± 7 seconds (Fig. 1i). Bacteria induced significantly more plasma membrane protrusions in cells where F-actin was depolymerized with cytochalasin D (CytoD) compared to control cells (Fig. 1j and k). This is in line with previous studies demonstrating that plasma membrane remodeling by meningococcus does not depend on actin polymerization^11,16,17^. Plasma membrane protrusions in CytoD-treated cells also showed significantly shorter lifetimes (Fig. 1l), further suggesting that F-actin stabilizes plasma membrane protrusions once they are formed and pointing to a role of membrane tension in this remodeling. Isogenic bacteria deficient in pilus retraction were still capable of inducing plasma membrane protrusions (*pilT* mutant, Fig. 1m and n), showing that forces exerted by pilus retraction were not required to remodel the host cell plasma membrane, as previously proposed for large bacterial aggregates^17^. Pilus retraction-deficient bacteria elicited discrete protrusions (Fig. 1n and Extended Data Fig. 2), as well as ring-like structures (Fig. 1m and o), reflecting more protrusions that are not resolved optically and larger protrusions (Extended Data Fig. 2) likely due to the higer amount of T4P produced by the *pilT* mutant. Strikingly, discrete membrane protrusions induced by *pilT* were no longer dynamic. Once they were formed, they never disappeared during the time of observation (Fig. 1p), suggesting a causal link between pilus retraction and disappearance of plasma membrane protrusions. When taken together, these observations prompted us to hypothesize that plasma membrane protrusions are induced by a direct interaction of the plasma membrane with T4P fibers.

**Figure 2.**
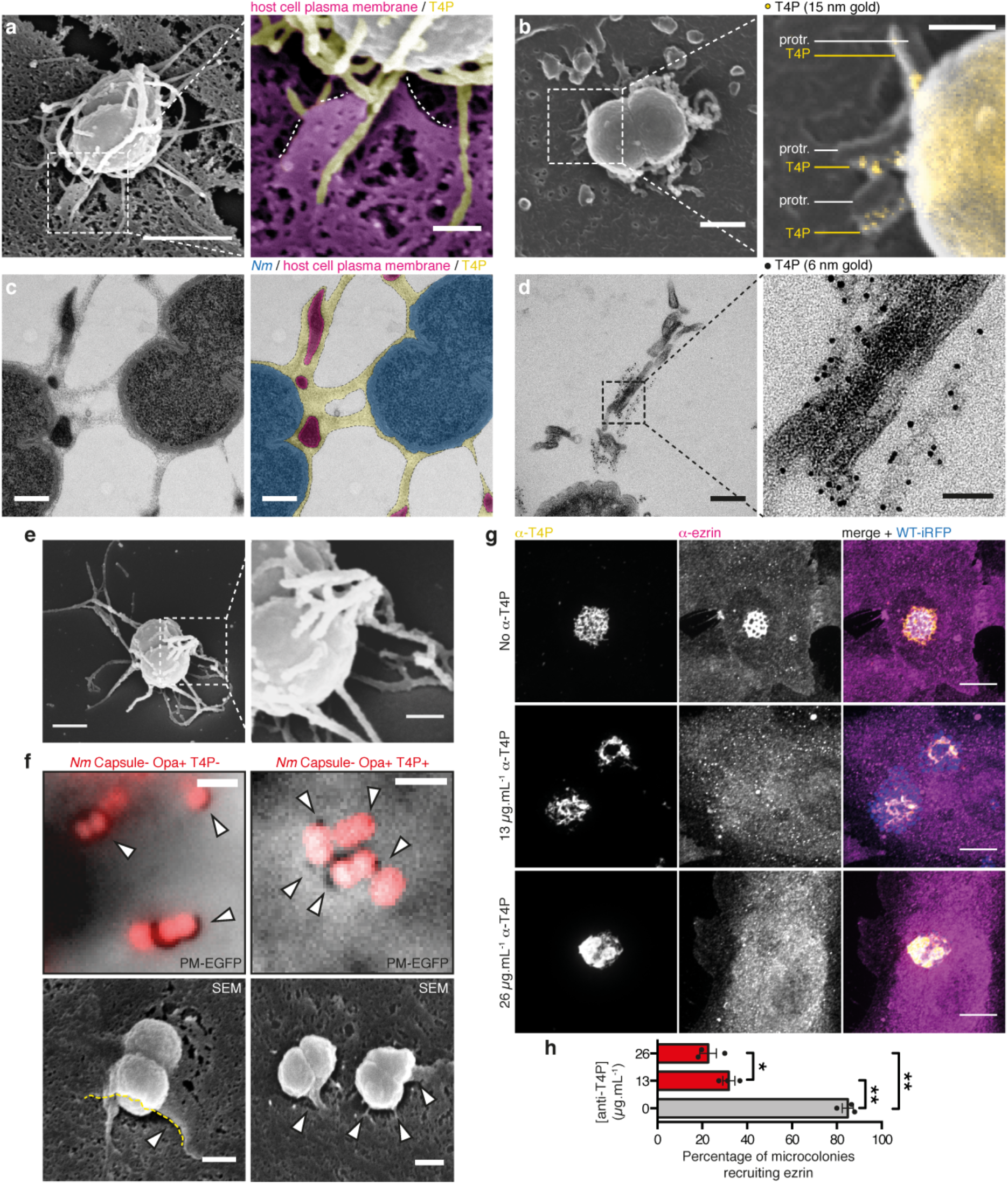
The morphology of plasma membrane protrusions is dictated by the fiber-like morphology of T4P. **(a)** Scanning electron micrograph of an endothelial cell after 10 min infection and stabilization of the T4P with a monoclonal antibody. Colorized crop shows plasma membrane protrusions (magenta, dotted lined) attached alongside T4P fibers (yellow). Scale bars, 1 μm and 200 nm. n=2 independent experiments performed in duplicate. **(b)** Same experiment after immunogold labeling of T4P. Protr., protrusions. Scale bars, 500 nm and 200 nm. n=2 independent experiments performed in duplicate. **(c)** Transmission electronmicrograph of a microcolony of *Nm* after high pressure freezing and freeze substitution, showing plasma membrane protrusions embedded in a meshwork of T4P. Scale bars, 200 nm. Representative of multiple microcolonies in n=1 experiment. **(d)** Same experiment after immunogold labeling of T4P. Scale bars, 200 nm and 50 nm. Representative of multiple microcolonies in n=1 experiment. **(e)** Scanning electron micrograph of the meshwork of T4P produced by an individual bacterium in culture. Scale bars, 500 nm and 200 nm. n=2 experiment. **(f)** Non-piliated Opa+ bacteria remodel the plasma membrane in a phagocytic cup-like fashion. Re-expression of T4P reverts the remodeling to a protrusion morphology. Scale bars, 2 μm (darkfield) and 500 nm (SEM). n=3 and 1 experiments. **(g)** An anti-T4P moclonal antibody affects plasma membrane remodeling by *Nm*. Plasma membrane recruitment was assessed by the accumulation of ezrin. Scale bars, 10 μm. Representative of n=3 experiments. **(h)** Quantification of the experiment in **(g)**. Bars and error bars represent the mean ±SEM. Dots represent the mean of individual experiments. 216 to 240 microcolonies were counted per condition.

As a first evidence, 3D Structured Illumination Microscopy showed that plasma membrane protrusions were intertwined with T4P fibers within pairs of meningococci (Extended Data Fig. 3a and b) and larger bacterial aggregates (Extended Data Fig. 3c and d). Next, to gain ultrastructural information, we performed scanning electron microscopy after stabilization of T4P, which are otherwise fragile, by coating them with a monoclonal antibody. In these conditions, numerous fiber-shaped structures were visible (Fig. 2a, compare with Fig. 1b). These fibers stained positive for T4P in immunogold labeling assays (Fig. 2b) and showed direct association with plasma membrane protrusions (Fig. 2a and b). While the majority of T4P fibers formed a dense meshwork around bacterial bodies (Extended Data Fig. 4a), some other extended further away from bacteria and were as long as 20 μm (Extended Data Fig. 4b). However, plasma membrane protrusions were only seen associated with T4P in the vicinity of bacterial bodies, and not with T4P fibers adhering flat to the plasma membrane (Extended Data Fig. 4c). At longer times of infection, 2 hours post-infectionn, high pressure freezing and freeze substitution prior to transmission electron microscopy revealed that plasma membrane protrusions were embedded in a dense meshwork-like structure that occupied the space between adjacent bacteria in the microcolony (Fig. 2c and Extended Data Fig. 5a). This meshwork was sometimes present around bacteria with no neighboring bacterium and still contained plasma membrane protrusions (Extended Data Fig. 5b). Immunogold labeling of T4P confirmed that this meshwork was made of T4P fibers (Fig. 2d). It should be emphasized that 2 hours after the initial contacts between pili and host cells bacteria have proliferated, the cells have moved and reorganized their cytoskeleton thus generating a complex picture that has been visualized in *Neisseria gonorrhoeae^21^*. Nevertheless, these data point to a scaffolding mechanism for plasma membrane remodeling exerted by T4P fibers. Several lines of evidence support this hypothesis. First, scanning electron microscopy of bacterial cultures showed that the T4P meshwork pre-existed adhesion with host cells (Fig. 2e). Second, adhesion of non-piliated bacteria to endothelial cells mediated by a non-fibrillar outer membrane adhesin, Opa, and its cellular receptor, CEACAM1, led to a phagocytic cup-shaped remodeling of the plasma membrane instead of discrete protrusions (Fig. 2f). SEM images show bacteria nearly engulfed in plasma membrane. Furthermore, re-expression of T4P reverts plasma membrane reorganization to dot-like protrusions, demonstrating a geometrical effect of T4P fibers on the shape of plasma membrane remodeling (Fig. 2f). Both membrane remodeling processes were again independent of F-actin polymerization (Extended Data Fig. 6). Finally, presence of an anti-T4P antibody during the infection of endothelial cells strongly impaired the ability of bacterial microcolonies to remodel the plasma membrane in a dose-dependent manner (Fig. 2g and h), suggesting that plasma membrane remodeling by meningococcus depends on the adhesive properties of T4P fibers. Therefore, we propose that the remodeling of the plasma membrane is driven by adhesion forces between receptors in the membrane bilayer and T4P fibers. The pilin monomer which is the main constituent of TFP was shown to interact with a heterodimer formed by CD147 and the β2-adrenergic receptor^22^.

We hypothesized that membrane remodeling occurs by membrane wetting along T4P fibers and explored the physical processes involved. It is known that a liposome spreads onto micrometer-scale adhesive glass fibers^23^. In this case, the free energy of the phospholipid bilayer spreading on a fiber of radius *r* is the sum of three contributions: (i) the curvature energy of the bilayer, (ii) the surface energy of the bilayer and (iii) the gain of adhesion energy between the bilayer and the fiber: 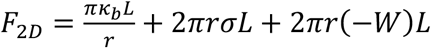, where *κ_b_* and *σ* are the bending modulus and surface tension of the bilayer, respectively, *L* is the spreading length of the membrane along the fiber, and Wis the adhesion energy of binding of mobile receptors on the membrane to ligands on the fiber. From *F*_2*D*_, we derived the driving force of the spreading of the bilayer, 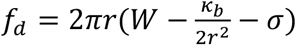. For the bilayer to spread, *f_d_* has to be positive. Therefore, *f_d_* = 0 defines a minimal value of the radius 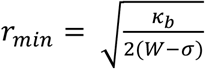. In conclusion, the bilayer spreads and wraps around the fiber, a wetting process that is already well known, but only when *f_d_* > 0, i.e. when *r > r_min_* (Fig. 3a).

**Figure 3.**
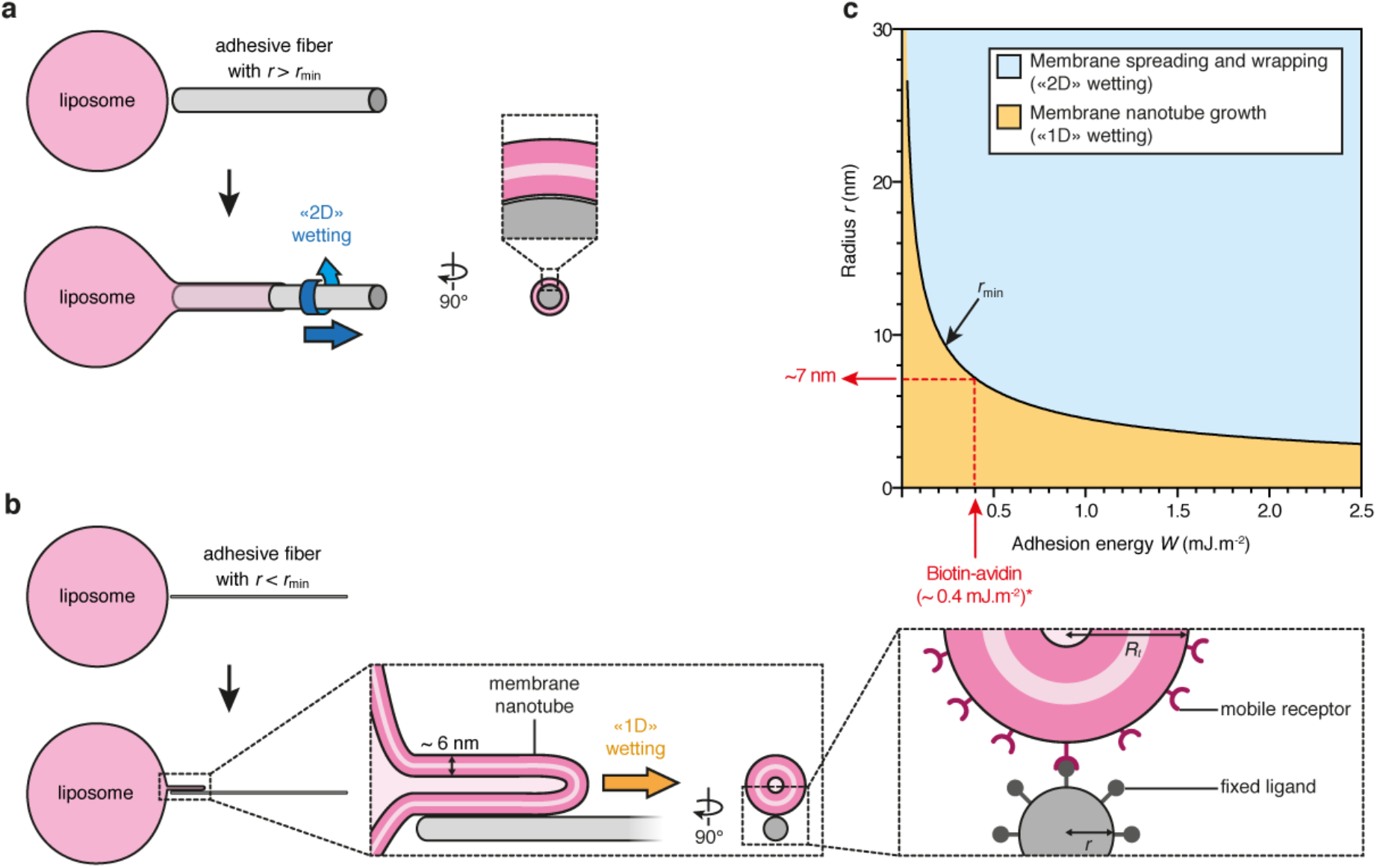
Theoretical prediction of a novel regime of wetting for the spreading of a liposome on an adhesive nanofiber. **(a)** A liposome spreads on a fiber of radius *r > r_min_* by “ 2D” membrane wetting, where the bilayer wraps around the fiber. **(b)** On a fiber of radius *r < r*_min_, the liposome cannot wrap the fiber because of the curvature energy of the bilayer and is predicted to grow a tube of radius *R_t_* along the fiber, which we call “ 1D” membrane wetting. In the inset most to the right mobile receptors and fixed ligands are indicated schematically, their size or distribution are not drawn to scale. (c) Phase diagram of a membrane bilayer spreading on a fiber versus nanofiber radius *r* and adhesion energy *W*. The black line corresponds to *r_min_* and separates the “ 2D” (blue region) and “ 1D” (yellow region) membrane wetting regimes. (*) Considering a biotinylated actin filament decorated with NeutrAvidin and a molar ratio of biotinylated actin:actin of 1:10 (Fig. 4), we estimated the adhesion energy *W* for the biotin-avidin complex to be 0.4 mJ.m^-2^, yielding a minimum radius *r_min_* = 7 nm below which “ 2D” wetting can no longer occur (See Materials and Methods).

The case of *r < r_min_*, which would correspond to spreading on a nanofiber, was not reported before. In this case, the bilayer cannot wrap around the fiber because the curvature energy becomes too high and a different scenario is predicted by the model: a membrane nanotube grows on top of the fiber, provided that the membrane tension is low enough (Fig. 3b). In this case, the free energy *F*_1*D*_ of a membrane tube of radius *R_t_* and length *L_t_* growing along an adhesive fiber is the sum of the tube energy and the gain of the adhesion energy 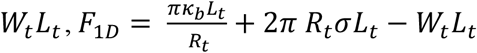. Here, the two first terms are the membrane curvature energy and the surface energy; *W_t_* = Г_*l*_ · *U*, where *U* is the ligand-receptor binding energy and Г_*l*_ is the number of ligands on the fiber per unit length. From the free energy *F*_1*D*_, we derived the driving force acting on the membrane nanotube, 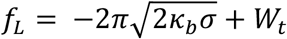 (See Materials and Methods). Here, the first term is the retraction force acting on the tube while the second term is the driving force due to the adhesion along the fiber. If *f_L_* > 0, *i.e*. when the surface tension *σ* of the membrane is sufficiently low, a membrane nanotube can grow along the fiber. To summarize, our model predicts that, for a given adhesion energy between receptors in the phospholipid bilayer and an adhesive fiber, there is a minimum radius *r_min_* of the fiber above which the bilayer should spread by canonical membrane wetting (“2D”) on the fiber, and below which the bilayer should spread by forming a nanotube along the fiber (“1D”). This last regime of tube formation is very different from other mechanisms of tube formation where tubes are actively pulled by molecular motors^24,25^ or by external mechanical forces^26-29^. For 1D wetting, the driving force is provided by the gain of adhesion energy. We show in Figure 3c the phase diagram of membrane nanofiber wetting in the coordinates *r* (fiber radius) and *W* (adhesion energy). The two regimes “2D” versus “1D” are separated by the curve 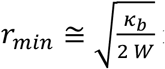 in the limit *σ ≪ W*. The dynamics of protrusions for 1D and 2D wetting is ruled by the balance between the driving force discussed above and the drag force associated to the flow of the lipids (See Materials and Methods). T4P fibers with their extremely small radius of 3 nm can thus potentially function in the “ 1D” membrane wetting regime. Indeed, in the context of live cells where membrane rigidity is higher (*κ_b_* ≈ 50 *k_B_T*^30^), the adhesion energy necessary for “2D” wetting, defined by *r_min_ = r_pilus_* = 3 *nm*, is extremely large, of order 10 mJ.m^-2^, which means that for T4P fibers we expect “1D” wetting to occur (Extended Data Fig. 7). This is supported by scanning electron micrographs showing plasma membrane protrusion along T4P fibers (Fig. 2a and b).

To test this theoretical model *in vitro* the use of purified pili and lipid vesicles was complicated by the necessity to insert a multimeric protein receptor in the lipid vesicles^22^. We rather used an assay consisting of model membranes (biotinylated giant unilamellar vesicles (GUVs) of typically 20 μm diameter) interacting with model nanofibers consisting of biotinylated actin fibers decorated with NeutrAvidin and immobilized on a solid substrate. The diameter of single actin filaments, 7 nm, is similar to the diameter of T4P fibers. Confocal microscopy showed that biotinylated GUVs interacting with NeutrAvidin-coated filaments deformed along them in two different ways. On thick adhesive bundles, GUVs spread and wrapped around the fibers (Fig. 4b), similar to previous studies of GUVs spreading on glass microfibers^23^ and consistent with the canonical “ 2D” membrane wetting described in our phase diagram (Fig. 3c). However, on thin adhesive nanofibers, GUVs tended to extend thin protrusions aligned along these fibers (Fig. 4c), consistent with the regime of membrane nanotube elongation that is predicted to occur on thin nanofibers (Fig. 3c). Cryo-electron microscopy experiments demonstrated that when phospholipid vesicles were mixed with adhesive nanofibers, membrane nanotubes were clearly in close apposition with the fibers rather than wrapped around them (Extended Data Fig. 8). Strikingly, membrane protrusions branched when encountering branched adhesive fibers (Fig. 4c), showing that the shape of the bilayer was governed by the architecture of the adhesive fibers. Interestingly, by taking reported values of the adhesion energy between biotin and avidin^31,32^, we found that the minimum radius *r_min_* below which the membrane should spread on these adhesive nanofibers by nanotube elongation is approximately 7 nm (Fig. 3c, and see Materials and Methods), in agreement with our experimental observations.

**Figure 4.**
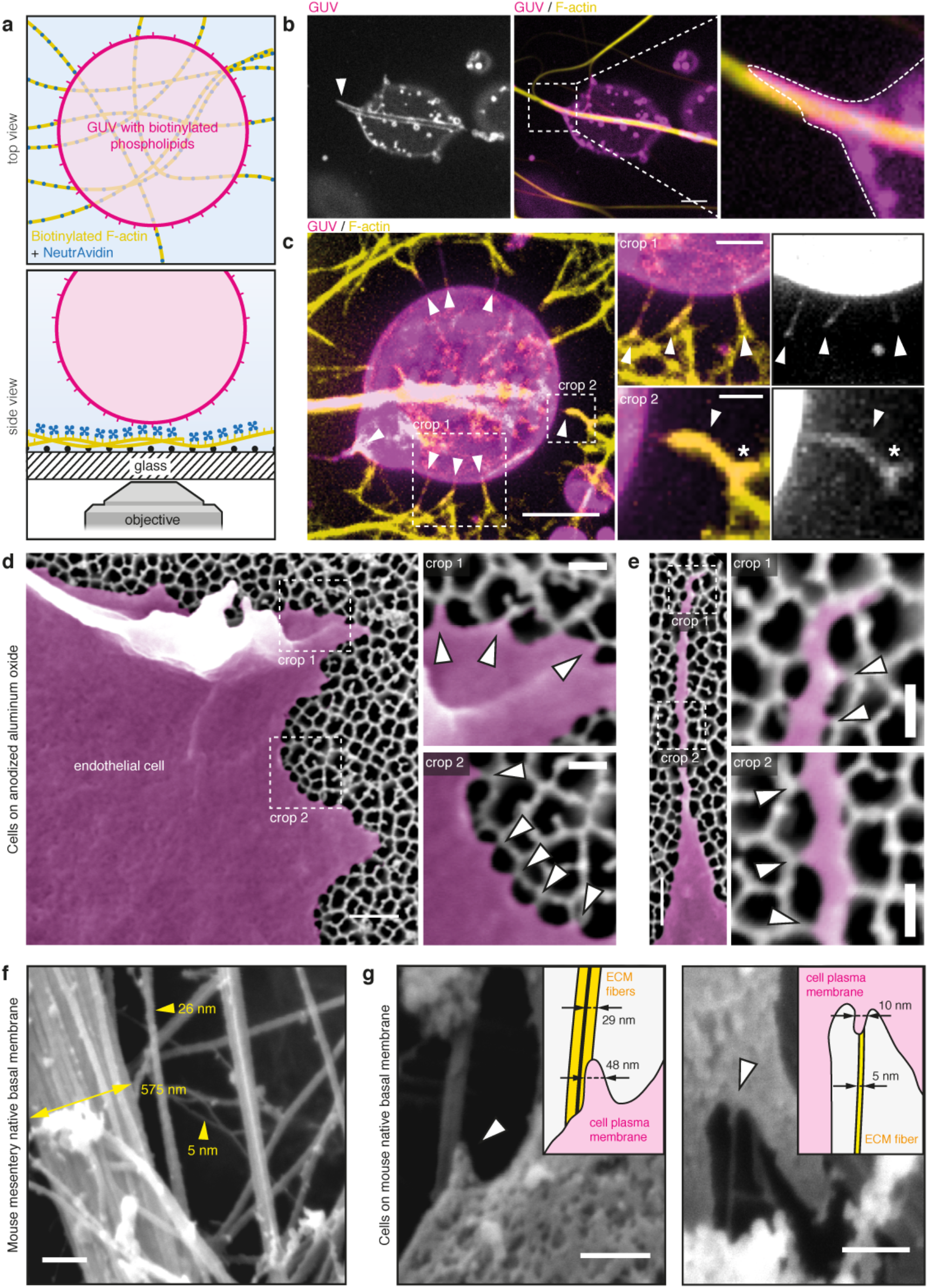
Membrane wetting on fibers drives the deformation of giant unilamellar vesicles and occurs in human endothelial cells. **(a)** Schematic of the experimental setup where giant unilamellar vesicles (GUVs) containing biotinylated phospholipids adhere to biotinylated F-actin fibers decorated with NeutrAvidin. **(b)** GUV s spread on thick adhesive bundles (arrowhead) by wrapping around the fibers (crop, dotted line), as predicted by the canonical “ 2D” membrane wetting regime. Scale bar, 5 μm. **(c)** On thin adhesive fibers, GUVs extend small membrane tubes that align with the fibers (arrowheads), as predicted in the “ 1D” membrane wetting regime. Membrane tubes can branch when encountering branched adhesive fibers (crop 2, asterisk). Scale bars, 5 μm, 2 μm and 1 μm. Representative of n=3 experiments. **(d)** Endothelial cells cultured on porous aluminum filters coated with an RGD peptide show local linear plasma membrane deformations that match with the thin pore walls (arrowheads). **(e)** Pore walls dictate the path followed by filopodia, where the plasma membrane is also found to deform locally when encountering a pore wall (arrowheads). Representative of n=2 experiments. Scale bars, 500 nm and 150 nm for the crops. **(f)** SEM shows that native basal membranes from mouse mesentery feature fibers with diameters down to 5 nm. Scale bar, 250 nm. **(g)** Endothelial cells cultured on such extracellular matrices show nanoscale plasma membrane protrusions aligning on nanoscale fibers. Scale bars, 250 and 200 nm.

Interaction with the extracellular matrix is another situation in which cells interact with nanofibers. To further validate our predictions in the context of the interaction of human cells with components of the extracellular matrix, we investigated how the plasma membrane of endothelial cells deforms on an RGD peptide-coated nanostructured solid substrate. To do so, endothelial cells were cultured on anodized aluminum oxide foils featuring 100 nm pores with pore walls of less than 15 nm mimicking fibers of the extracellular matrix (Fig. 4d and e). To reconstitute integrin-fibronectin adhesion without altering the nano-topography of the substrate, we sequentially coated the substrate with an azide functionalized PLL-PEG (azido-(poly-L-lysine-graft-poly[ethylene glycol], or APP) and a reactive RGD peptide (BCN-RGD)^33^. Scanning electron microscopy showed that the plasma membrane at the cell contour deformed periodically along the pore walls (Fig. 4d). Filopodia were also seen to follow the complex geometry of the pore walls along their whole length (Fig. 4e). Interestingly, the plasma membrane along filopodia also seemed to deform periodically when encountering a pore wall. To test if protrusions formed on natural ECM nanofibers we performed scanning electron microscopy on native basal membranes extracted from mouse mesentery (Fig. 4f). The extracellular matrices exhibited bundles of fibers of a few tens or hundreds of nanometers, but also fibers down to a few nanometers. When cultured on such matrices, human endothelial cells showed nanoscale plasma membrane protrusions that aligned onto fibers of 5 to about 30 nm (Fig. 4g). Thus, this set of experiments shows that cell membranes can form protrusions when adhering on naturally occurring nanofibers such as those found in the ECM.

Taken together, these data show that adhesive nanofibers can drive the remodeling of biological membranes at the nanoscale because of a wetting phenomenon that falls into a new regime that we named “ 1D” membrane wetting. In the context of plasma membrane remodeling induced by *N. meningitidis* in human endothelial cells, we propose that adhesion to T4P fibers drives plasma membrane deformation by this mechanism (Extended Data Fig. 9). This is consistent both with the fact that the process is F-actin independent^11,16,17^, and with the observation that actin polymerization occurs subsequent to membrane deformation^17^. A similar sequence is observed in membrane nanotube pulling experiments in which actin polymerizes in membrane nanotubes initially devoid of F-actin^34,35^.

Calculations based on our model indicate that the adhesion energy between T4P and the cell membrane should be smaller than 10 mJ.m^-2^ and the membrane tension should be smaller than a threshold value (*σ_t_*) corresponding to *f_L_* = 0 to be in the “ 1D” regime. Assuming pilin monomers are arranged in a helical pitch of 7 nm in the pilus fiber^36^, an estimation of the binding energy per pair of pilin monomer-cell receptor leads to a minimal value of approximately 100 *k_B_T* for “ 2D” wetting to occur. As a comparison, it can be estimated from atomic force microscopy measurements on *Pseudomonas aeruginosa* T4P that the binding energy is smaller, approximately 18 *k_B_T* (see Materials and Methods), which leads to an estimate of the driving force *W_t_* in the order of 10 pN. Thus, membrane wetting on T4P is likely to enter the “ 1D” regime, provided that the tube retracting force is smaller than *W_t_*. The retraction forces of tubes have been measured for different cell types adhering on substrates with various techniques and are in the range of a few to 20 pN^37-39^. Available estimates of key physical parameters are thus in agreement with the occurrence of “ 1D” wetting during *N. meningitidis* interaction with host cells.

Membrane deformation as discrete protrusions along T4P fibers could be seen as a way to prevent the formation of a continuous membrane cup, as seen for the zipper mechanism employed by *Listeria monocytogenes* or *Yersinia* spp. were receptor containing plasma membrane adheres to outer membrane invasins^40,41^, leading to bacterial engulfment. Therefore, “ 1D” wetting along filamentous adhesins could be a strategy to prevent efficient internalization into host cells. Of note, many prokaryotic species, including numerous human pathogens, also produce filamentous appendages which organize into dense meshworks of fibers^42-53^ and which could possibly use a “1D” membrane wetting mechanism in the interaction with their environment or their immobilization on host cells.

Membrane nanotubes were already described to play important roles in other physiological as well as pathological cellular processes^54-58^. Our study therefore describes a new mechanism of membrane nanotubes formation that is central to the interaction between an infectious bacterium and its host cell, but also to normal cell physiology. Indeed, in eukaryotic multicellular organisms, cells interact with proteins of the extracellular matrix that exist as fibers with adhesive properties, such as collagen, fibrillin or fibronectin, with reported diameters as small as 10 nm^59^. Our data show that fibers of the extracellular matrix can be even smaller, down to 5 nm. Importantly, cells form plasma membrane protrusions on such nanofiber networks. Thus, it will be of interest to assess whether such an F-actin-independent “ 1D” membrane wetting phenomenon plays a role in cell migration through complex environments occurring during diverse biological processes such as development, immune response or cancer.

Our study highlights the notion that cells react to the topography of their environment at the nanoscale^60,61^. It is likely that the development of high resolution light and electron imaging techniques will help uncover the cell response to nano-topographical cues, in particular in the context of the interaction with complex fibrous environments.

## Acknowledgements

This work was supported by the French ministry for research and higher education (ACO); the Integrative Biology of Emerging Infectious Diseases (IBEID) laboratory of excellence (GD) and the VIP European Research Council starting grant (GD). We gratefully acknowledge the Imagopole – Citech of Institut Pasteur (Paris, France) as well as the France– BioImaging infrastructure network supported by the French National Research Agency (ANR-10–INSB–04; Investments for the Future), and the Région Ile-de-France (program Domaine d’ Intérêt Majeur-Malinf) for the use of the Zeiss LSM 780 Elyra PS1 microscope and the Zeiss Auriga scanning electron microscope. The authors greatly acknowledge the Cell and Tissue Imaging (PICT-IBiSA), Institut Curie, member of the French National Research Infrastructure France-BioImaging (ANR10-INBS-04), and the PICT-IBiSA Institut Curie (Paris, France). This work was supported by Institut Curie, Fondation pour la Recherche Médicale (FRM FDT20170437130) and Ecole Doctorale Frontières du Vivant (FdV)– Programme Bettencourt (RS), ERC StG grant STARLIN (DMV), Centre National de la Recherche Scientifique (CNRS), and P. B. and F.B.-W. belong to the CNRS consortium CellTiss, to the Labex CelTisPhyBio (ANR-11-LABX0038) and to Paris Sciences et Lettres (ANR-10-IDEX-0001-02). F.C.T. was funded by the EMBO Long-Term fellowship (ALTF 1527-2014) and Marie Curie actions (H2020-MSCA-IF-2014, project membrane-ezrin-actin). We thank J. Pernier for discussions, providing actin and advice on actin reconstitution. We thank Rafaele Attia and Jian (Olivier) Shi for technical help. We would like to thank Sven van Teeffelen for discussions and for help with the quantitative analysis of dark field microscopy. We would like to thank Dorian Obino for critical reading of the manuscript. The authors declare no competing financial interests.

## Author contributions

A.C.-O. designed, conducted and analyzed most of the experiments, and prepared all the figures. G.D. designed the experiments, analyzed the data and supervised the project. F.-C.T. designed and performed the GUVs on adhesive actin fibers experiments. F.-C.T., F.B.-W. and P.B. performed the mathematical analysis of membrane wetting. D.B. performed and analyzed the micropatterns experiments. V.M. and A.C.-O. performed and analyzed the fluorescence experiments in mice. M.S. and A.C.-O. performed and analyzed the HPF-FS TEM experiments. A.M. and A.C.-O. conducted and analyzed the SEM experiments. A.S. and A.C.-O. performed and analyzed the 3D-SIM experiments. K.M. performed the TEM experiment in the mouse. R. S. prepared the mouse basal membranes. F.-C.T. and A.B. performed the cryo-EM experiments. C.M. constructed the pMGC13 plasmid. S.G. and P.L. produced the nanobody against T4P used in the mice experiments. M.P. contributed to the design of the Anodisc experiments and to the interpretation of the data. A.C.-O., F.-C.T., B.-W., P.B. and G.D. wrote the manuscript. All the authors edited the manuscript.

---

**Extended Data Figure 1.**
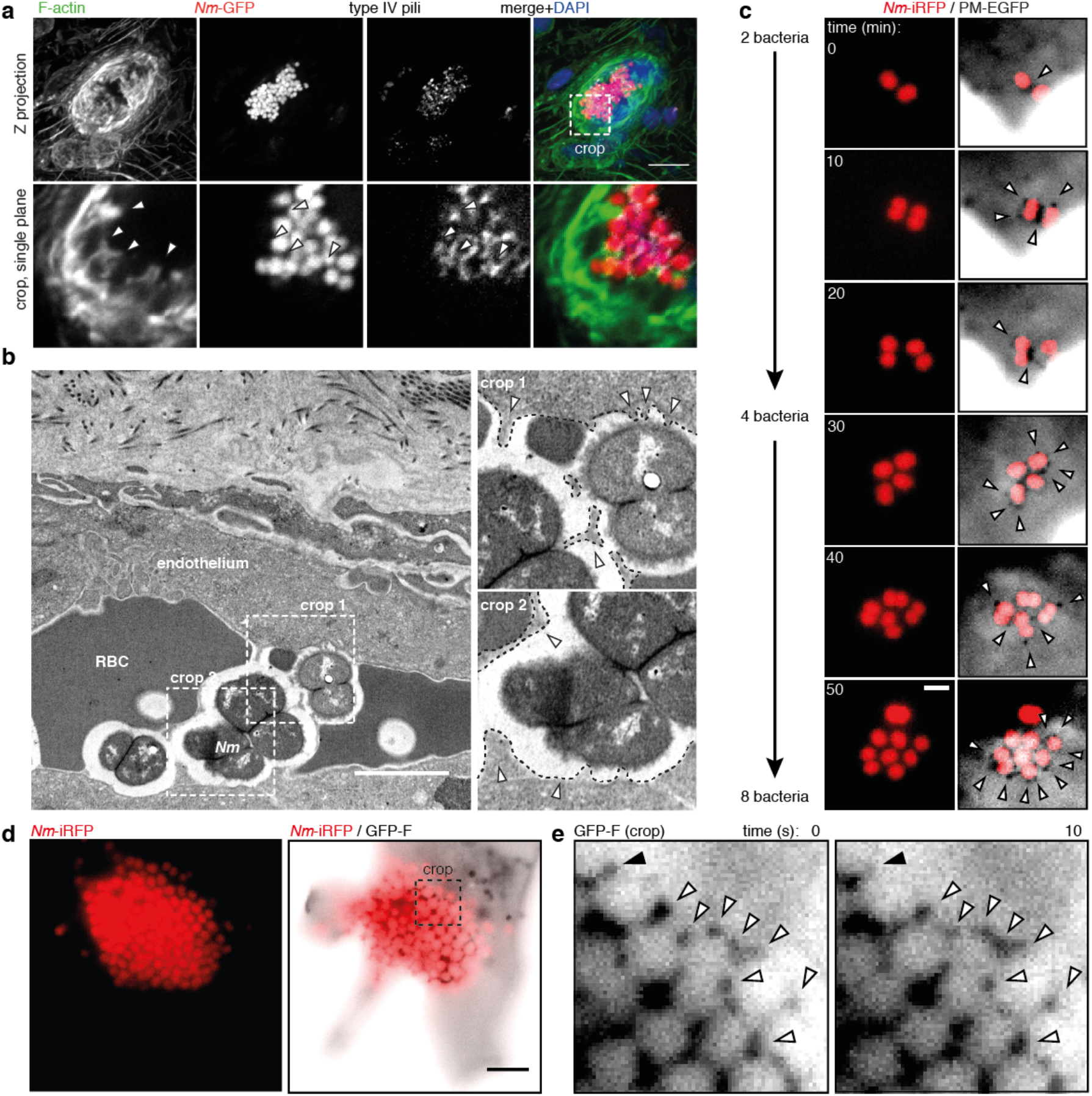
Further characterization of plasma membrane protrusions induced by *Nm in vivo* and by individual bacteria *in vitro*. **(a)** Histoimmunolabeling of human blood vessels in a mouse after 3h of infection with *N. meningitidis*(*Nm*)-GFP showing plasma membrane protrusions containing F-actin (arrowheads). Scale bar, 10 μm. Representative of n=2 mice, **(b)** Transmission electron micrograph showing the plasma membrane of endothelial cells (dashed lines in the crops) remodeled beneath and between aggregated bacteria in an infected human vessel (arrowheads). Scale bar, 2 μm. n=1. **(c)** Oblique illumination live imaging of a micropatterned endothelial cell expressing the plasma membrane marker PalmitoylMyristoyl-EGFP (PM-EGFP, inverted contrast) infected by individual Λfa/–iRFP. Plasma membrane protrusions initiated at the level of two bacteria are accumulated as bacteria divide on the host cell surface and remain within the nascent microcolony (arrowheads). Scale bar, 2 μm. Representative ofn=3 experiments. (d-e) Oblique illumination live imaging of an endothelial cell expressing the membrane marker GFP-F infected by an aggregate of Nm-iRFP shows that plasma membrane protrusions in this case are no longer dynamic (white arrowheads) with rare events of disappearing protrusions occurring at the edge of the bacterial aggregate (black arrowhead). Representative of n=3 experiments. Scale bar, 5 μm.

**Extended Data Figure 2.**
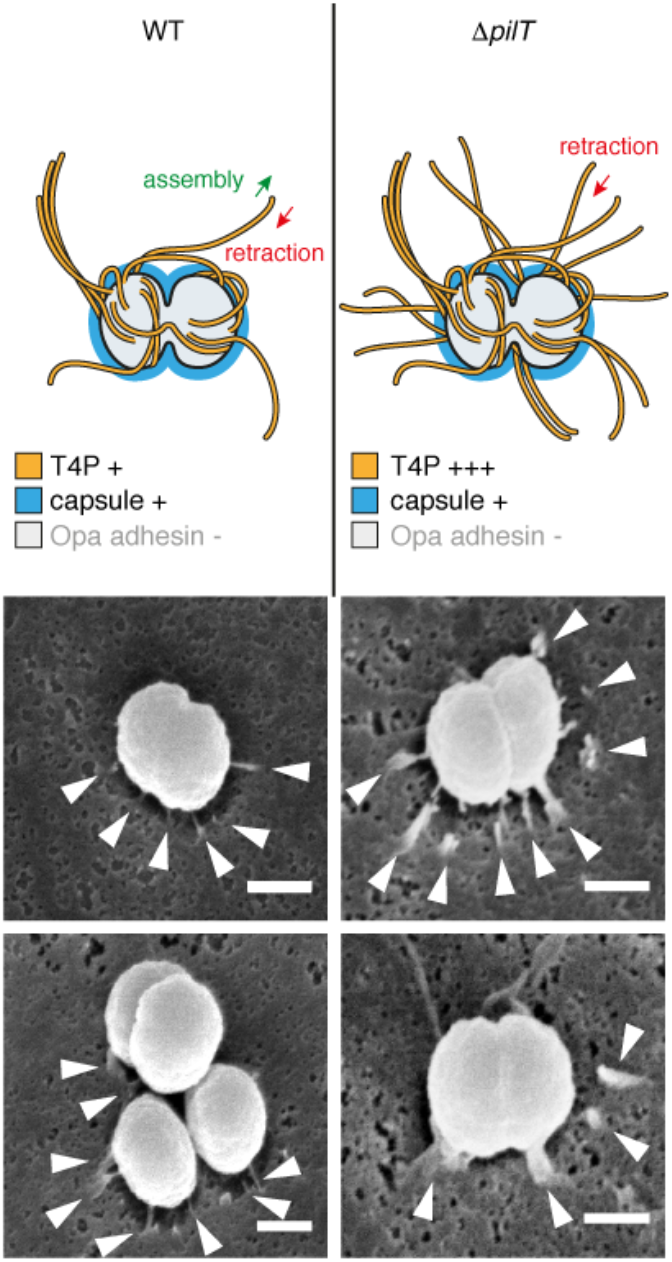
T4P retraction-deficient individual bacteria trigger plasma membrane protrusions. Scanning electron microscopy showing that the *pilT* mutant, which cannot retract T4P, still induces plasma membrane remodeling in the form of discrete protrusions as in the wild-type strain (WT). However, either more protrusions or protrusion slightly larger are observed, likely due to the higher amount of T4P produced by the *pilT* strain. Note that T4P are not visible in this preparation. Scale bars, 500 nm. n=1 experiment.

**Extended Data Figure 3.**
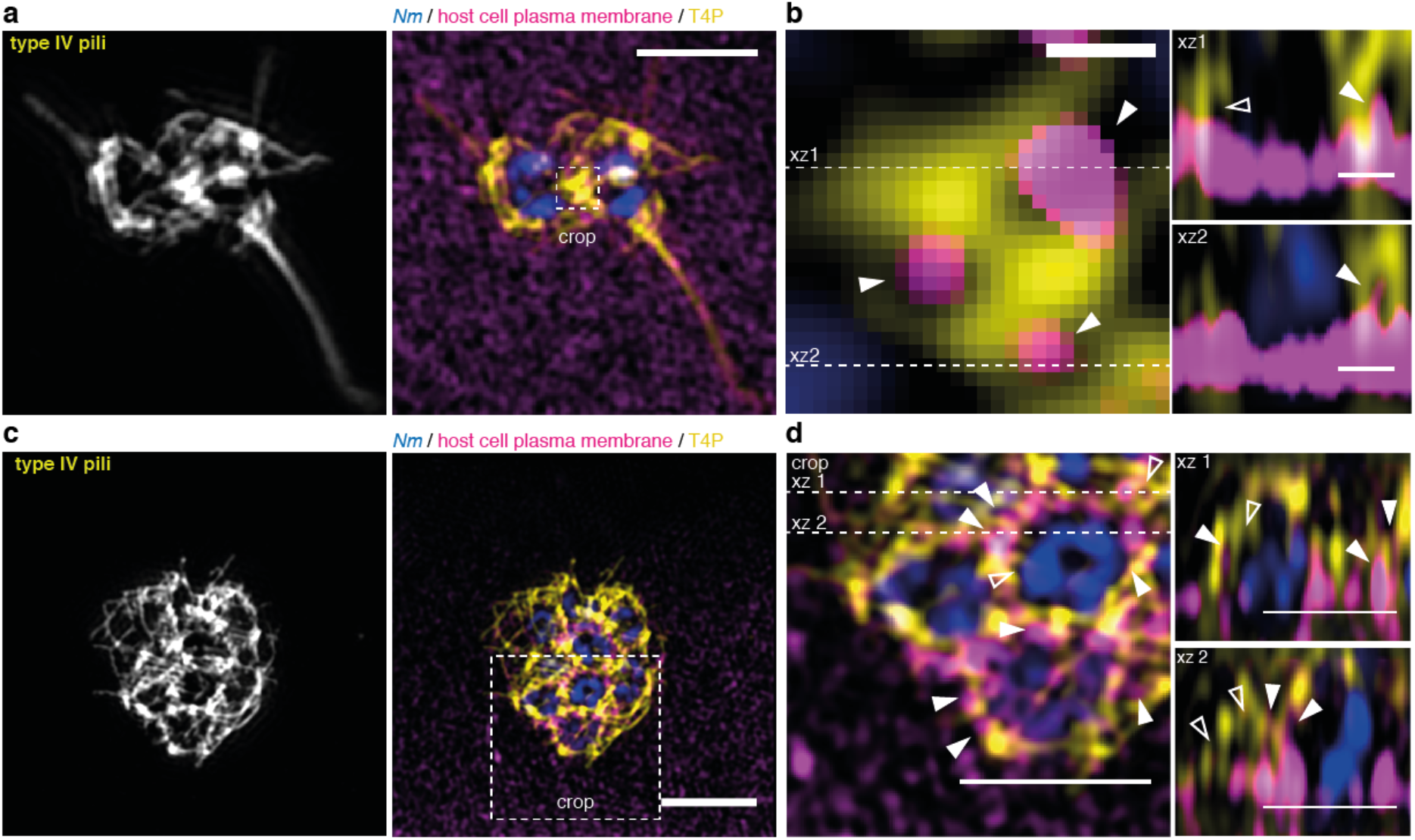
T4P-plasma membrane interface visualized by 3D SIM. **(a)** 3D Structure Illumination Microscopy (SIM) micrographs of a pair **(a, b)** and a small aggregate of 8 **(c, d)** meningococci (DAPI, blue) after 30 min infection of endothelial cells expressing the membrane marker PM-EGFP and immunostained for T4P and GFP. **(a)** and **(c)** show T4P detection and merged images in Z projections. Scale bars, 2 μm and 3 μm. **(b)** and **(d)** are cropped merged images and Z-sections showing details of T4P organization with empty spaces between fibers (empty arrowheads) and spaces occupied by plasma membrane protrusions (filled arrowheads). Scale bars in **(b)**, 200 nm for the first inset then 500 nm. Scale bars in **(d)**, 2 μm. n=3 experiments.

**Extended Data Figure 4.**
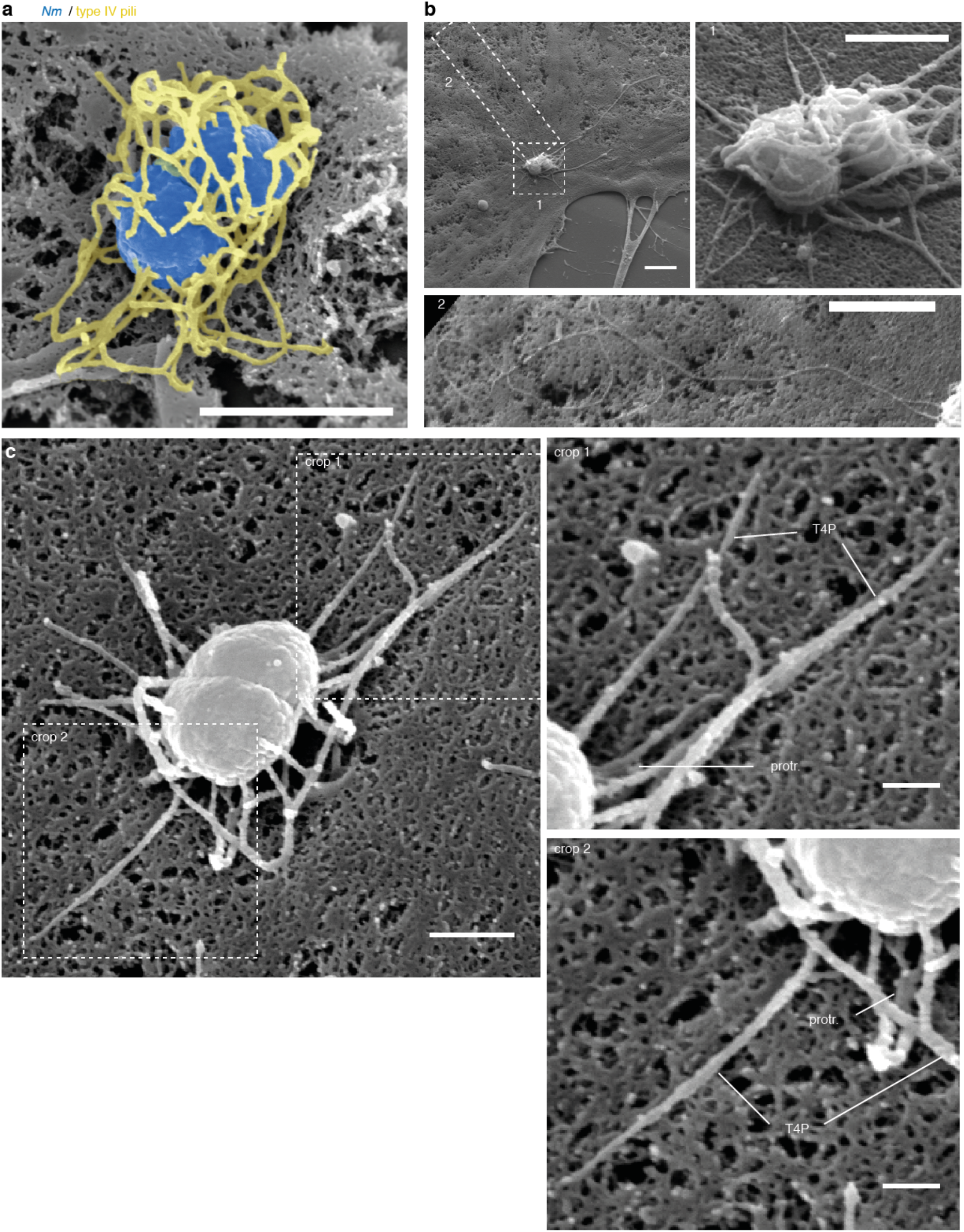
Additional examples of *Nm* T4P architecture and T4P-plasma membrane interface by SEM after stabilization of the T4P with a monoclonal anti-T4P. **(a)** Example of a single bacterium (false colored in blue) featuring a very dense meshwork of T4P (false colored in yellow) that encloses the bacterial body. Scale bar, 1 μm. **(b)** Example of a pair of bacteria (crop 1) with particularly long T4P (crop 2). Scale bars, 10 μm (large view) and 2 μm (crops). (c) Example of a single bacterium where host cell plasma membrane forms protrusions near the bacterial body but not along T4P fiber away from the bacterium, as better seen in the crops. Protr., protrusions. Scale bars, 500 nm (whole picture) and 200 nm (crops). n=2 experiments.

**Extended Data Figure 5.**
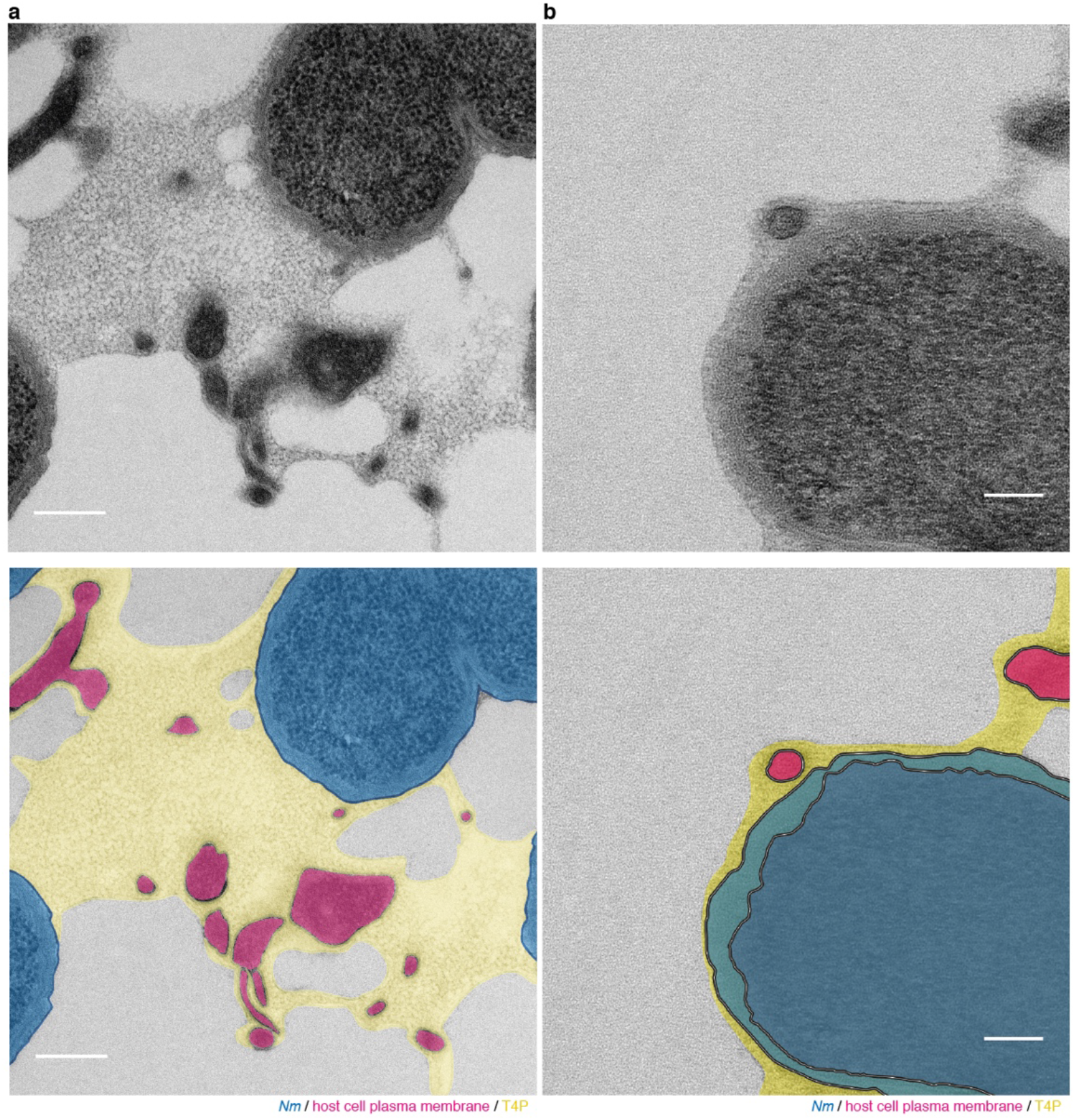
Additional examples of the T4P-plasma membrane interface visualized by TEM after HPS-FS. Transmission electron micrographs and colorized micrographs of microcolonies of *Nm* on endothelial cells after 2h of infection, high pressure freezing and freeze substitution, **(a)** Example of a dense meshwork of T4P embedding plasma membrane protrusions. Scale bar, 200 nm. **(b)** Example of a protrusion that lies in a layer of T4P at the periphery of a bacterium with no neighboring bacteria. Scale bar, 100 nm. Representative of multiple microcolonies in n=1 experiment.

**Extended Data Figure 6.**
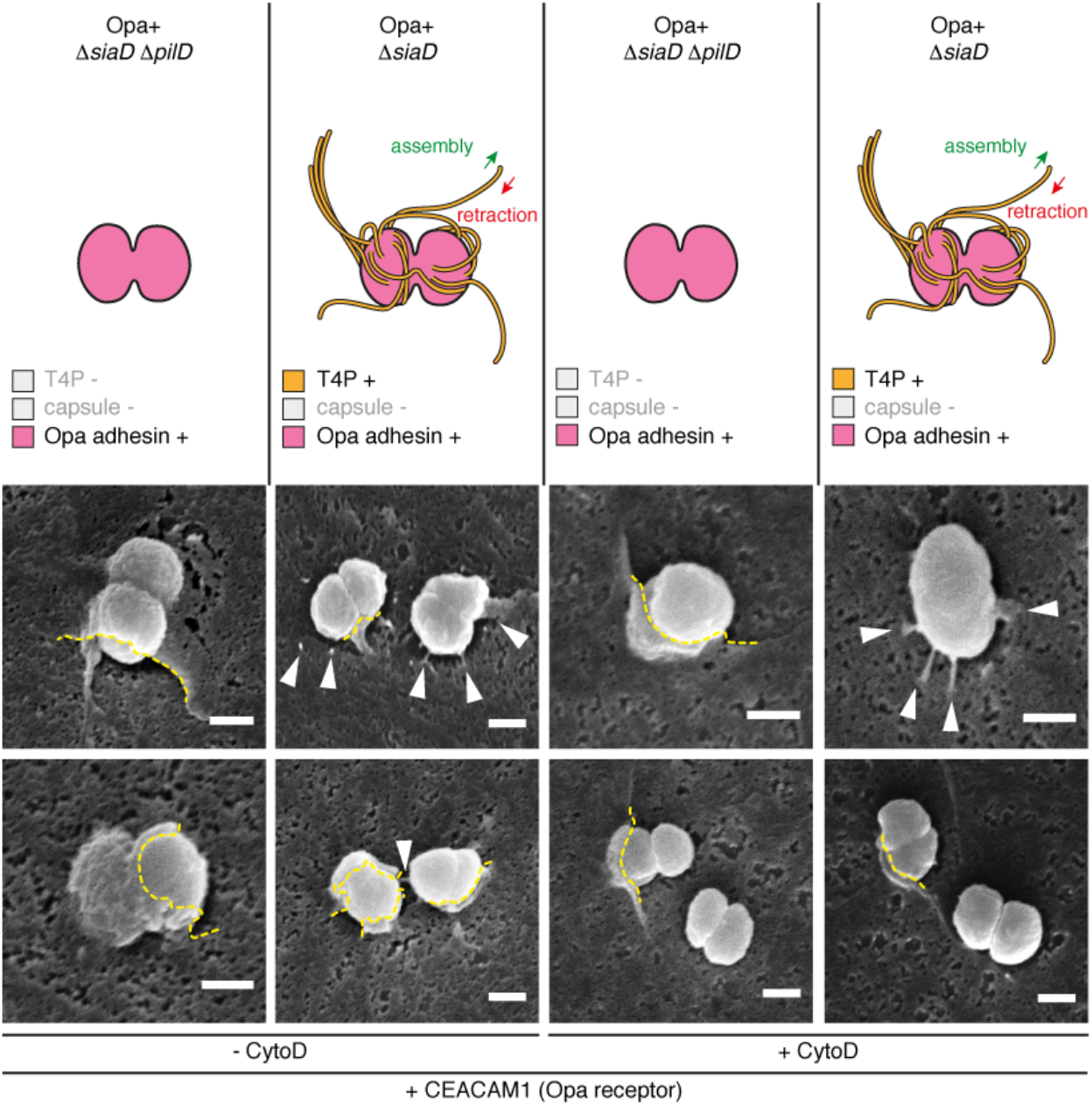
Adhesion via non-fibrillar adhesins leads to plasma membrane remodeling as a cup-like structure. Bacteria expressing the alternative Opa outer membrane adhesion elicit plasma membrane remodeling in a cup-like fashion in cells expressing the Opa receptor CEACAM1. Reexpression of T4P in the same genetic background leads to plasma membrane remodeling as a mix of discrete protrusions and incomplete cups. Depolymerization of the F-actin cytoskeleton with cytochalasin D (CytoD) prior to bacterial adhesion does not inhibit plasma membrane remodeling driven by adhesion to either Opa or T4P adhesins. Scale bars, 500 nm. n=1 experiment.

**Extended Data Figure 7.**
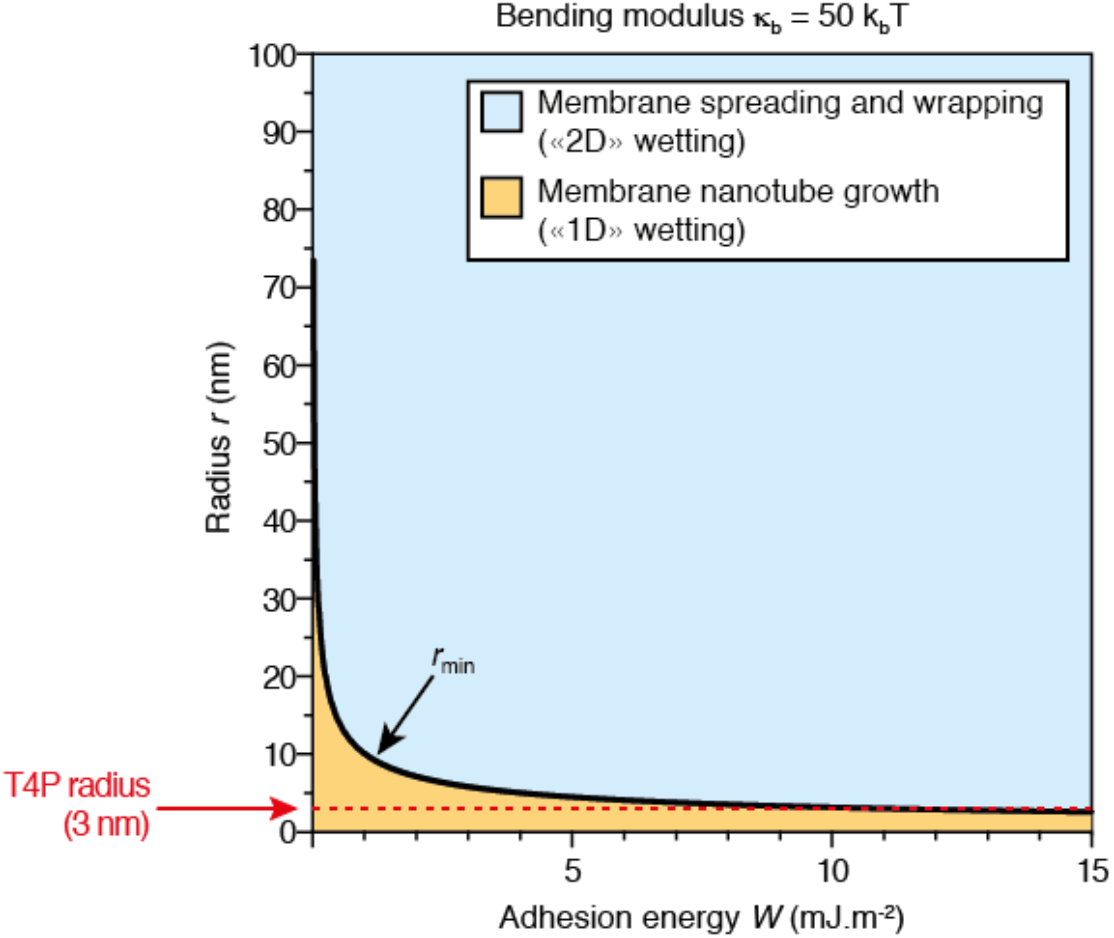
Theoretical prediction of “ 1D” and “ 2D” wetting regimes for the spreading of a cell membrane on an adhesive nanofiber. Phase diagram of a membrane bilayer spreading on a fiber versus nanofiber radius *r* and adhesion energy *W*. The black line corresponds to *r_min_* and separates the “ 2D” (blue region) and “ 1D” (yellow region) membrane wetting regimes. Here, the bending modulus *κ_b_ ≈* 50*k_B_T* is the one found in live cells_30_. The radius of a T4P fiber yields a theoretical minimal adhesion energy of 10 mJ.m_-2_ for “ 2D” cell membrane wetting to occur.

**Extended Data Figure 8.**
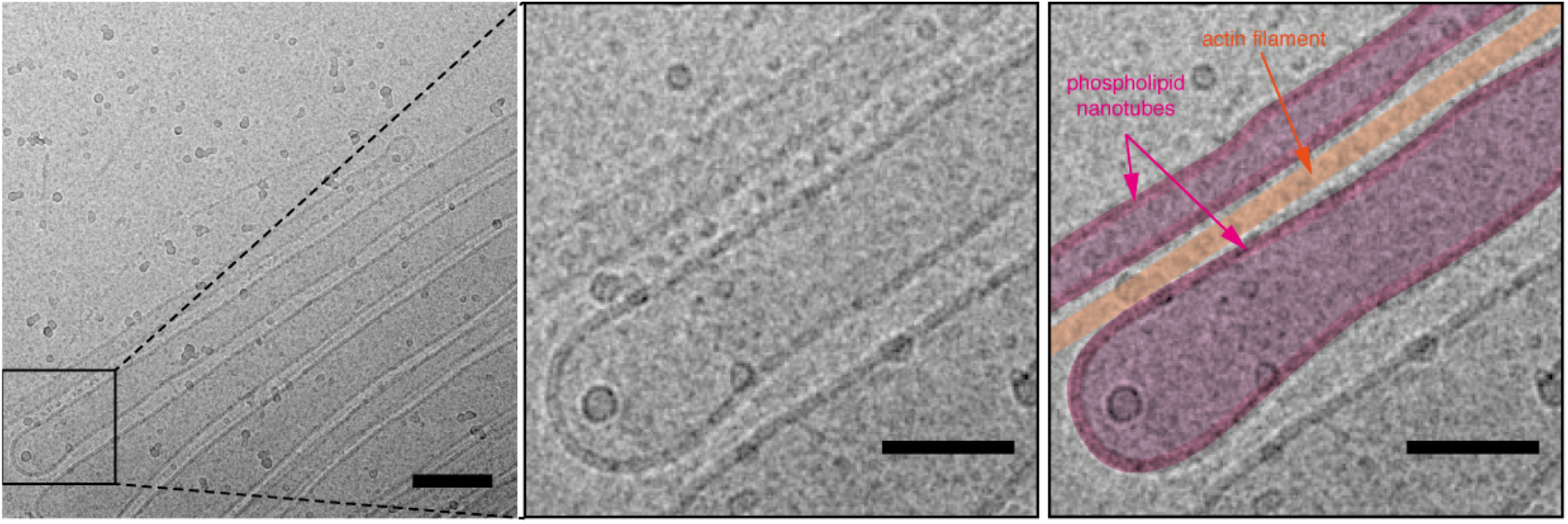
Cryo-electron microscopy of vesicles on nanofibers experiment. Here vesicles containing biotinylated phospholipids were mixed with biotinylated F-actin fibers decorated with NeutrAvidin, and imaged by cryo-EM (left). Membrane nanotubes were visibly aligned along individual F-actin filaments instead of wrapping around them (middle and right panels). Scale bars, 100 nm (left) and 50 nm (middle and right panels).

**Extended Data Figure 9.**
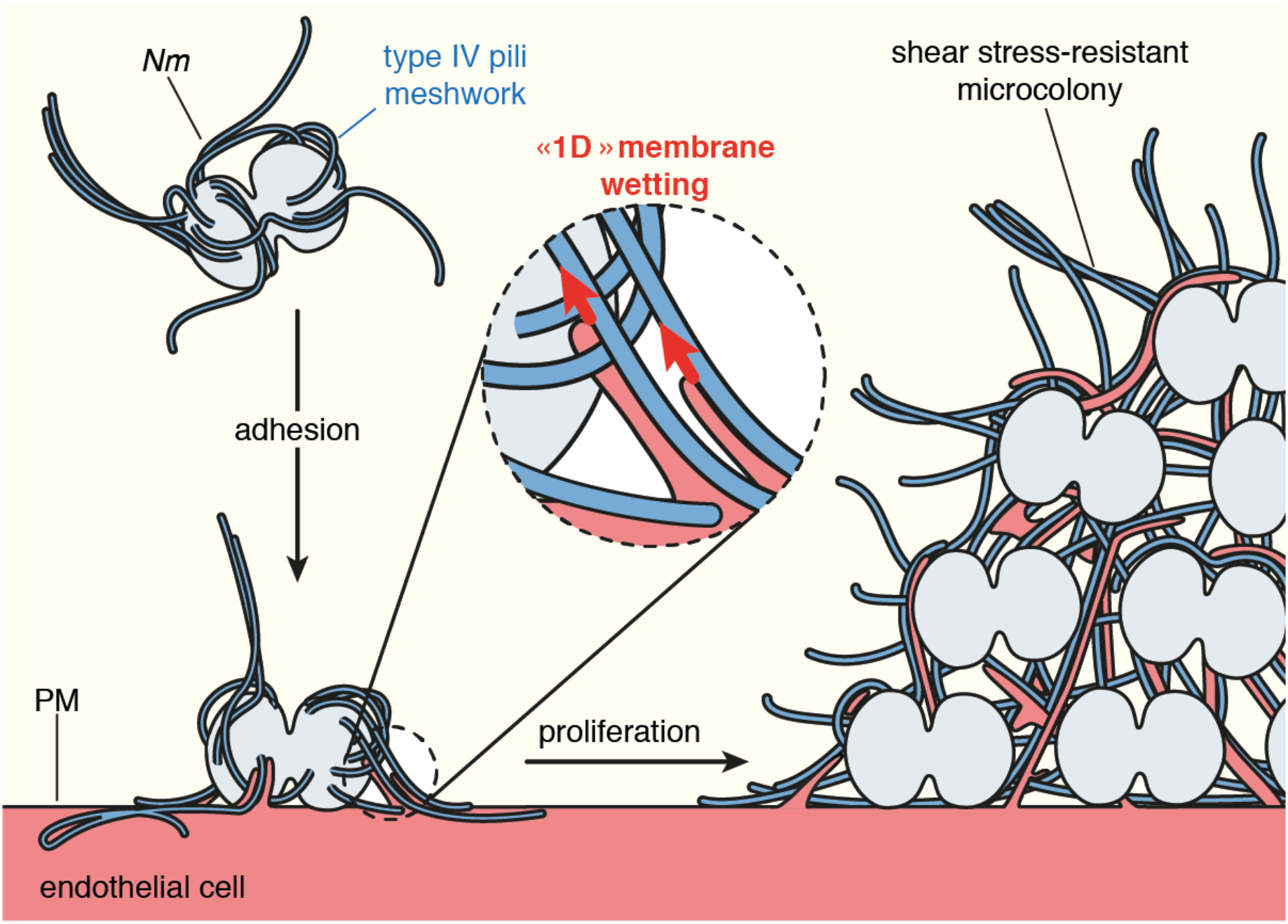
Working model for the mechanism of wetting-induced plasma membrane protrusions by *Nm* T4P fibers. *Nm* produces T4P as a meshwork of fibers. Upon adhesion to the host endothelial cell, the high adhesiveness of T4P allows “ 1D” membrane wetting of the plasma membrane, and thus remodeling of the membrane alongside T4P fibers. As the bacteria proliferate and aggregate extracellularly, plasma membrane protrusions remain attached to T4P fibers and end up embedded in a dense extracellular T4P meshwork. This complex T4P-plasma membrane structure provides the microcolony with enough mechanical coherence to resist blood flow-generated shear stress.

## Materials and Methods

### Ethics statement

All experimental procedures involving animals were conducted in accordance with guidelines established by the French and European regulations for the care and use of laboratory animals (Décrets 87–848, 2001–464, 2001–486 and 2001–131 and European Directive 2010/63/UE) and approved by the local ethical committee Comité d’ Ethique en matière d?xpérimentation Animale, Universite Paris Descartes, Paris, France. No: CEEA34.GD.002.11. All surgery was performed under anesthesia, and all efforts were made to minimize suffering. For human skin, written informed consent was obtained and all procedures were performed according to French national guidelines and approved by the local ethical committee, Comité d’ Evaluation Ethique de l’ INSERM IRB 00003888 FWA 00005881, Paris, France Opinion: 11—048.

### Animals, grafting and infection

Humanized SCID/Beige (SOPF/CB17 SCID BEIGE. CB17.Cg-Prkdc-Lyst/Crl) mice (Charles River, France) 5—8 weeks of age were used as described in earlier studies (Melican et al., 2013). Briefly, anesthetized mice were grafted with healthy human skin, including epidermis, dermis and dermal microvasculature. 3-4 weeks post graft, the tissue retained human morphology without inflammation. 10^7^ CFU of GFP expressing bacteria were injected IV and the animals were sacrificed 2h post infection.

### Preparation of tissue samples for immunohistochemistry

Tissues were fixed with 4% paraformaldehyde (PFA), frozen in OCT (Tissuetek) and sliced at 10 μm. Human vessels were stained for plasma membrane with the human specific *Ulex europaeus* agglutinin (UEA) lectin coupled to rhodamine (Vector Laboratories) and for F-actin with phalloidin coupled to AlexaFluor-568 (Invitrogen). Type IV pili were detected with an in-house generated anti-PilE nanobody coupled to AlexaFluor-647 (AlexaFluor-647-NHS-ester, Invitrogen).

### Preparation of tissue samples for transmission electron microscopy

Tissues were fixed with 4% PFA and 0.5% glutaraldehyde in 0.1M phosphate buffer, for 4h at room temperature. They were then transferred to 1% PFA in 0.1M phosphate buffer and incubated overnight at 4°C. Samples were next processed as for conventional TEM.

### Cell culture

Human umbilical vein endothelial cells (HUVEC, Promocell) were maintained in Endo-SFM (human endothelial-SFM, Gibco) supplemented with 10% (v/v) heat-inactivated fetal bovine serum (FBS, PAA Laboratories) containing 10 μg.mL^-1^ endothelial cells growth supplement (ECGS, Alfa Aesar) and were used between passages 3 and 9.

### Bacterial strains and culture

*Neisseria meningitidis* (Nm) serogroup C strain 8013 is a capsulated, piliated, Opa^-^, Opc^-^ clinical isolate (Nassif et al., 1993). It was grown on GCB-Agar (Difco) plates containing Kellogg’ s supplements (Kellogg et al., 1968) and antibiotics when required (kanamycin, 100 μg.mL^-1^; chloramphenicol, 5 μg.mL^-1^; spectinomycin, 50 μg.mL^-1^; erythromycin, 2 μg.mL^-1^) at 37°C and 5% CO2 under moist atmosphere. For infection experiments, GCB-Agar grown bacteria were resuspended in Endo-SFM supplemented with 10% FBS at an OD600 of 0.05 and cultured for 2 h with shaking at 37°C and 5% CO2. The GFP expressing strain was described elsewhere (Melican et al. 2013). A plasmid allowing stable expression of the iRFP near-infrared fluorescent protein (Filonov et al., 2011) in *Nm*, pMGC13, was constructed as follows: the sequence encoding the iRFP protein was PCR-amplified from the plasmid pBAD/His-B-iRFP (a kind gift from Vladislav Verkhusha, Addgene plasmid #31855) with PacI and SalI restriction sites in 5’ and 3’ respectively and the ribosome binding site from *Nm pilE* gene just upstream of the start codon. The sequences for the primers are iRFP_F2: <u>TTAATTA</u>*AAGGAGTAATTTTATGGGGGGTTCTCATCATCATCA* and iRFP_R3: <u>GTCGAC</u>TCACTCTTCCATCACGCCGATCTGC (with restriction sites underlined and RBS from *pilE* in italic). The PCR fragment was then cloned in a pCRII-TOPO vector (Invitrogen), checked for sequence and subcloned in the pMGC5 plasmid that allows homologous recombination at an intergenic locus of the Nm chromosome and expression under the control of the constitutive *pilE* promoter (Soyer et al., 2014). The pilT-iRFP mutant strain deficient for type IV pili retraction was generated by homologous recombination of the chromosomal DNA of the *pilT* insertion mutant strain into the wild-type iRFP strain by natural transformation. The non-capsulated Opa+-siaD-iRFP and its non-piliated derivative Opa+-siaD-pilD-iRFP was generated by homologous recombination of the chromosomal DNA from *siaD* and/or *pilD* insertion mutant strains into a naturally occurring variant of 2C43, which carries a *opaB* gene in the ON phase, by natural transformation.

### Cell transfection and plasmids

5.10^5^ HUVEC were electroporated with 4 μg plasmid DNA encoding either the enhanced green fluorescent protein fused to palmitoylation and myristoylation signals from the Lyn protein (PM-EGFP, kindly provided by Barbara Baird and David Holowka, Cornell University, Ithaca), a farnesylated GFP (GFP-F, Clonetech), and/or LifeAct-mCherry (kindly provided by Guillaume Montagnac, Institut Curie, Paris) or the Carcinoembryonic antigen-related cell adhesion molecule 1 (CEACAM1, kindly provided by Alain Servin) with the Amaxa Nucleofector device (Amaxa Biosystem, Lonza).

### Oblique illumination (dark field) live cell imaging and high speed analysis

2,1.10^4^ transfected cells were seeded in 96-well glass bottom plates (Sensoplate, Greiner BioOne) coated with 3 μg.mL^-1^ rat tail type I collagen and 50 μg.mL^-1^ human fibronectin (Sigma-Aldrich) and cultivated without ECGS. The next day, cells were infected with vortexed Endo-SFM-FBS grown bacteria at a MOI of 200 and gently rinsed after 10 min with Endo-SFM-FBS and screened for presence of adhering bacteria prior imaging. For inhibition of F-actin polymerization, cells were treated with 100 nM cytochalasin D in Endo-SFM-FBS for 20 min at 37°C and 5% CO2 prior to infection. Live imaging was performed on an Eclipse T*i* inverted microscope (Nikon) equipped with a laser-based iLas2 Total Internal Reflection Fluorescence microscopy (TIRF) module (Roper Scentific) with lasers at 491 nm, 561 nm and 647 nm and an ORCA03 digital CCD camera (Hamamatsu). Dark field images were obtained by using the TIRF module to get the illumination lasers with an incident angle of 31.2° in the widefield mode through an oil immersion 100x Apo TIRF objective with a 1.49 numerical aperture and alternatively an additional 1.5x lens. A 2−2 binning was used. To perform spatial high-resolution acquisition, a fast Z-piezo stage (Piezo Nano Z100) was adapted onto the stage of the microscope. For imaging in the stream mode, images were recorded at the camera frame rate (5 to 6 images per second) for 10 seconds. Focus was maintained with the Perfect Focus System (PFS, Nikon). All experiments were performed at 37°C in an incubation chamber adapted for the microscope (Microscope Temperature Control System, LIS). The set-up was controlled by the MetaMorph software (Molecular Devices). Times of appearance and disappearance of the plasma membrane protrusions around each bacterium were analyzed manually. kon and koff were calculated by dividing the total number of appearing or disappearing protrusions in all the bacteria of a given experiment by the total number of protrusions observed and by 10 seconds, and were consequently expressed in s^-1^. The mean lifetime for plasma membrane protrusions was calculated as the inverse of koff and was therefore expressed in seconds.

### Oblique illumination live cell imaging of HUVEC on micropatterns

Micropatterning on glass coverslips was performed as previously described^62^. Briefly, after plasma activation (Plasma Cleaner, Harrick), we passivated the glass with PLL-g-PEG (Surface Solutions GmbH, 0,1 mg.mL^-1^ in 10 mM HEPES pH 7.4) for 30 min. After washing with water, we illuminated the surface with deep UV light (UVO Cleaner) through a chromium synthetic quartz photomask (Toppan). For imaging, we used channel-shaped sticky slides (Ibidi) that we bound on top of micropatterned coverslips. The home built chambers were sterilized by 10 minutes UV exposure under the hood. Finally, we incubated 1h with human fibronectin (50 μg.ml^-1^ in water) and seeded PM-EGFP expressing HUVEC cells at a density of 50,000 cells per channel. 1h post seeding, cells were rinsed 3 times with fresh Endo-SFM to get rid of non-adhering cells, and cultured overnight. For infection, iRFP and mCherry wild-type bacteria from liquid cultures were vortexed and loaded into the channels in a 1:10 ratio to a final MOI of 400. Samples were incubated for 10 minutes at 37°C and 5% CO2 and then rinsed thoroughly 3 times with fresh Endo-SFM to remove non-adhering bacteria. To follow membrane remodeling during bacterial proliferation, oblique imaging with 491, 561 and 642 nm lasers was performed with a 100X ApoTIRF objective (Z-stacks: 1 μm steps over 10 μm; time-lapse: 10 min time frame over 10h).

### Scanning Electron Microscopy (SEM)

1,5.10^5^ HUVEC infected with *Nm* for 10 min at a MOI of 400 or bacteria alone cultured on 12 mm glass slides were chemically pre-fixed by addition of one volume of pre-warmed 8% PFA in 0,1M HEPES pH 7.4 and incubated at room temperature for 45 min. After three washing steps in HEPES, type IV pili were immunostained or not after 20 min blacking in HEPES-0.2% gelatin (HEPES-G) with 1,2 μg.mL^-1^ 20D9 antibody and a fluorescent secondary antibody or 2,4 μg.mL^-1^ 20D9 antibody and a secondary antibody coupled to 15 nm colloidal gold particles (EM.GMHL15, BBI Solutions) diluted 1:30. HEPES was used instead of PBS in all steps. Infected cells were then post-fixed with 2.5% glutaraldehyde in HEPES, overnight at 4°C, washed in HEPES, post-fixed with 1% OsO4 in HEPES for 1h, washed in distilled water, dehydrated in graded series of ethanol (25, 50, 75, 90 and 100%), critical point dried with Leica EM CPD300, mounted on aluminum stubs and sputter-coated with 20 nm gold/palladium with a Gatan PEC 682 gun ionic evaporator. SEM was performed in an Auriga scanning microscope (Carl Zeiss) operated at 7kV with an in-lens secondary electrons detector. For immunogold labelled samples, images were simultaneously acquired through the secondary electrons detector and the backscattered electrons detector at 20kV. Colorized images were generated with Adobe Photoshop.

### High pressure freezing, freeze substitution and transmission electron microscopy (HPF-FS-TEM)

3−0.05 mm sapphire disks (M. Wohlwend GmbH) were cleaned in 100% ethanol, carbon coated in a Baltec MED 010 evaporator, baked at 130°C for 8h, glow discharged for 5 sec in a Harrick PDC-32G-2 air plasma cleaner at low radiofrequency and coated immediately with 2,5 μg.mL^-1^ rat tail type I collagen and 50 μg.mL^-1^ human fibronectin. Disks were seeded with 1,5.10^5^ cells in 24-well plates. The next day, cells were infected with *Nm* for 2h at a MOI of 400 and chemically pre-fixed by addition of one volume of a mix of pre-warmed 8% PFA and 1% EM-grade glutaraldehyde in 0,1 M HEPES pH 7.4 and incubated at room temperature for 45 min. Cells were rinsed three times in 0,1 M HEPES. Disks were then spaced by a golden O-ring, sandwiched between a 0.4 mm and a 0.5 mm copper spacer, high pressure frozen in a Baltec HPM 010 and stored in liquid nitrogen. Samples were freeze substituted in a RMC Boeckler FS-8500 with a mix of 1% OsO4, 0.1% uranyl acetate and 5% water in glass distilled acetone. Freeze substitution cycle was as follows: -90°C for 1h, 2.5°C increase per hour for 16h, -50°C for 30 min, 15°C increase per hour for 2h, -20°C for 30 min, 10°C increase per hour for 2h and 0°C for 1h. After four washes in acetone, sample were embedded in Epon (Epon 812, AGAR1031 kit, hard formula, Agar 100). Alternatively, type IV pili were immunogold labelled before embedding. In this case, cells were chemically prefixed with 4% PFA and 0.1% glutaraldehyde (final concentrations), rinsed three times in HEPES, quenched with 50 mM NH4Cl in HEPES for 5 min, blocked with HEPES-G for 20 min, incubated with 12 μg.mL^-1^ 20D9 antibody in HEPES-G for 1h, washed twice in HEPES-G, incubated with goat anti-mouse antibody coupled to 6 nm colloidal gold particles (EM.GMHL6, BBI Solutions) diluted 1:30 in HEPES-G for 30 min, washed five times in HEPES-G and post-fixed with 1% glutaraldehyde in HEPES for 15 min. Samples were then high pressure frozen and freeze substituted as described above. Ultrathin sections of 60 nm nominal thickness were cut with a Leica EM UC7 ultramicrotome and collected on carbon coated, formvar-supported hexagonal copper grids, contrasted with 4% uranyl acetate and 3% Reynold’ s lead citrate and imaged in a FEI Tecnai T12 transmission electron microscope operated at an accelerating voltage of 120 kV.

### Immunofluorescence on 2h infected cells

1,5.10^5^ HUVEC were cultured in 96-well glass bottom plates coated with 2,5 μg.mL^-1^ rat tail type I collagen and 50 μg.mL^-1^ human fibronectin and infected the next day with *Nm* at an MOI of 100. After 30 min adhesion in the presence of 0, 13 or 26 μg.mL^-1^ 20D9 antibody, unbound bacteria were washed three times with fresh cell culture medium and allowed to proliferate for 2h in the presence or not of the same concentrations of 20D9. Infected cells were then fixed with 4% PFA and processed for immunofluorescence as for 3D SIM except that the 20D9 staining was omitted in wells were the antibody was already present. Ezrin was detected with a rabbit polyclonal anti-ezrin antibody (kind gift of Paul Mangeat). Images were taken with the Nikon Ti Eclipse spinning disk through a 100X oil immersion objective and ezrin recruitment was detected manually.

### 3D Structured Illumination Microscopy (3D-SIM)

1,5.10^5^ PM-EGFP expressing cells were seeded onto 12 mm glass coverslips coated with 3 μg.mL^-1^ rat tail type I collagen and 50 μg.mL^-1^ human fibronectin and cultivated without ECGS. The next day, cells were infected for 30 min at a MOI of 200 with Endo-SFM-FBS grown bacteria and fixed by addition of one volume of pre-warmed 8% PFA in PBS and incubation at room temperature for 30 min. Cells were rinsed three times in PBS, permeabilized for 10 min with PBS-0.1% Triton-X100 and blocked for 30 min in PBS-0.2% gelatin (PBSG). Cells were then incubated for 1h with 2 μg.mL^-1^ rabbit polyclonal anti-GFP antibody to amplify the membrane GFP signal (A11122, Invitrogen) and 1,2 μg.mL^-1^ mouse monoclonal anti-PilE antibody 20D9 (Pujol et al., 1997) to detect type IV pili. Cells were incubated with 10 μg.mL^-1^ secondary goat anti-rabbit and goat anti-mouse polyclonal antibodies coupled to AlexaFluor-488 and -568 (Invitrogen) for 1h. DNA was detected with 0,5 μg.mL^-1^ DAPI. All antibodies and DAPI were diluted in PBSG. Coverslips were mounted in Vectashield mounting medium (Vector Laboratories) and sealed with clear nail polish. SIM was performed on the ELYRA system controlled by the Zen software (Carl Zeiss) equipped with diode lasers at 405 nm, 488 nm and 561 nm. Images were acquired with an oil immersion 100x Plan-Apochromat M27 objective with a 1.46 numerical aperture and an additional 1.6x lens. Nominal pixel size was 0.049 and 0.084 μm in the XY and XZ planes, respectively, using a 1−1 binning. Structured illumination grid pattern was shifted by five different phases and three different angles along the whole Z-stack. Super resolved images were computationally reconstructed and channels were aligned with 100 nm TetraSpeck beads (Invitrogen) mounted in Vectashield. To further avoid reconstruction artefacts, raw images were checked for bleaching using the SIMcheck plugin^63^ in Fiji^64^. Only data sets with total intensity variations below 20% over the whole Z-stack were used. We could detect however the presence of putative “ hammerstroke” artifacts difficult to avoid in SIM^65^. Contrast was linearly enhanced in Fiji for clarity.

### Mathematical description of membrane spreading (“ 2D” membrane wetting) and membrane nanotube growth (“ 1D” membrane wetting) along adhesive fibers

The free energy of the membrane spreading on a cylindrical fiber is the sum of three contributions: curvature energy, surface energy and gain of the binding energy between the adhesion molecules on the membrane and binders on the fiber,

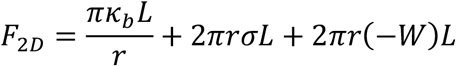

where *r* is the radius of the actin fiber, *κ_b_* is the bending modulus, *σ* is the membrane tension of the GUV, *L* is the spreading length of the membrane along the fiber, and *W* is the adhesion energy per unit area described as

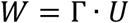

In this equation, *Γ* is the number of binders per unit area on the fiber and *U* is the adhesion energy per pair of adhesion molecule-binder.

From the free energy, we derived the driving force of membrane spreading, 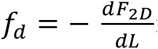;

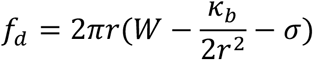

For membrane spreading to occur, *f_d_* has to be positive, leading to a minimum radius for an adhesive fiber

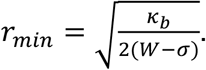

On a fiber of radius *r > r_min_*, the floppy vesicle spreads. As *L* increases, the tension of the vesicle increases because the excess membrane area decreases. Thus, the driving force *f_d_* decreases and the spreading stops when 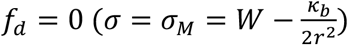.

For an adhesive fiber of radius *r < r_min_*, the vesicle cannot wrap around it. Instead, a membrane tube grows along the fiber. The free energy of a tubular membrane having a radius *R_t_* growing along the adhesive fiber with a length *L_t_* is the sum of the tube energy *F_t_* and the gain of the adhesion energy *W_t_L_t_*

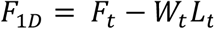

The tube energy is the sum of the curvature energy and the surface energy,

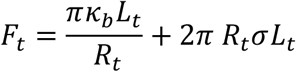

and *W_t_* is the adhesion energy of an adhesion molecule-binder pair per unit length, given by

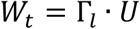

where Γ*_l_* is the number of bonds per unit length. *F*_1*D*_ depends upon two independent variables, *L_t_* and the volume 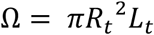. The force is coupled to *L*_t_, and the pressure inside the tube to *Ω*^66^. The force is classically derived from the total energy *F*_1*D*_ with respect to *L_t_* at constant volume, 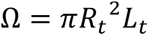, and is given by 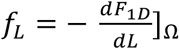. It leads to

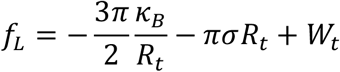

and the pressure inside the tube *p_t_* by the derivative of the free energy with respect to *Ω* at constant length; moreover, *p_t_* calculated from *F_t_* has to be equal to the pressure inside the vesicle, which is approximated to zero^29^. It leads to

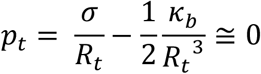

These two equations lead to

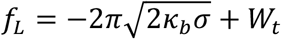

The first term is the retraction force acting on the tube^26-29, 67^ and the second term is the driving force pulling the tube.

### Numerical evaluation of *r_min_* in the case of GUVs spreading on biotinylated actin fibers

Given that the size of a NeutrAvidin molecule is 5 nm^68^, which is comparable to the diameter of an actin molecule, 4-7 nm^69^, and the molar ratio of biotinylated actin:actin is 1:10, Γ ≈ 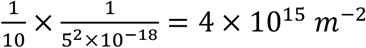. Taking 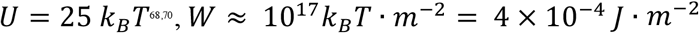. Here, for a floppy vesicle, taking 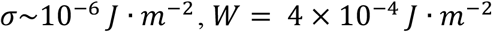 and *κ_b_* we obtained *r_min_* ≈ 7 *nm*. Assuming a cylindrical actin bundle composed of cylindrical actin filaments, we estimated that there should be at least 4 – 10 actin filaments in the actin bundle for membrane spreading to occur by “2D” wetting.

For a biotinylated actin fiber of radius *r < r_min_*, given the molar ratio of biotinylated actin:actin to be 1:10 and taking into account the helical structure of actin filaments, we estimate 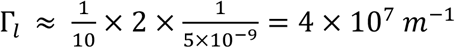. Taking 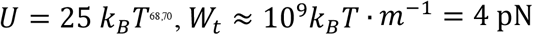. Taking value 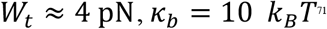, we find that 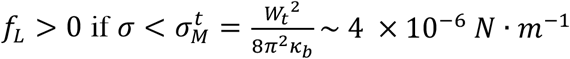.

### Dynamics of the two regimes

*2D wetting*. The dynamic of membrane spreading on the actin fiber with a radius *r* is characterized by 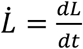 which can be derived from the balance between the driving force *f_d_* and the friction force *f_s_*.

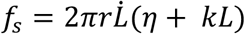

where *η* is a lipid viscosity and 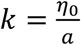 is friction coefficient, *η*_0_ is a molecular viscosity and *a* is a molecular length^23^. The friction force *f_s_* is the sum of friction due to the defect at the contact line between the vesicle and the spreading film, and the friction between the spreading membrane and the surface of the fiber. If the first term is dominant, the balance of the driving and friction forces leads to 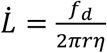. If the second term is dominant, the growth is diffusive at short time and 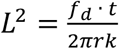.

*1D wetting*. For membrane nanotube growing along the actin fiber, 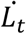 is determined by the balance between the driving force *f_L_* and the friction force *f_v_* introduced by E. Evans^26^

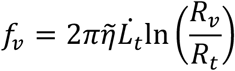

where 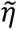 is a membrane viscosity, *R_v_* is the radius of the vesicle from where the membrane nanotube with a radius of *R_t_* grows. The origin of this force is due to the slip between the two monolayers at the entry of the tube: due to the cylindrical geometry, the flux of the lipids in the outer leaflet of the tube is larger than that in the inner one^26^. For short tubes, assuming *σ* is small, the balance of the driving force and the friction force leads to

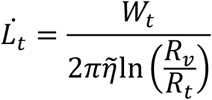

From this equation, taking from Figure 1f, 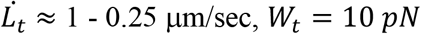 and 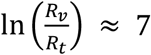, we obtained for the membrane surface viscosity 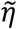 ranging between *2 × 10^−7^* and *10^−6^ N.sec.m^−1^* (Surface Poiseuille, SP), which are of the same order as the surface viscosities measured by R.E.Waugh for vesicles^72^.

### Interaction of giant unilamellar vesicles (GUVs) with adhesive nanofibers

*Reagents*. 1,2-dioleoyl-sn-glycero-3-phosphocholine (DOPC, 850375) and 1,2-distearoyl-sn-glycero-3-phosphoethanolamine-N [biotinyl(polyethyleneglycol)-2000] (DSPE-PEG(2000)-biotin, 880129P) were purchased from Avanti Polar Lipids. BODIPY-TR-C5-ceramide, (BODIPY TR ceramide, D7540) was purchased from Invitrogen. Hellmanex®II was purchased from Hellma (9-307-010-3-507). Culture-Inserts 2 Well for self-insertion were purchased from ibidi (Silicon open chambers, 80209). Biotinylated actin was purchased from Cytoskeleton (biotin-actin, AB07). Phalloidin-FluoProbes®488 was purchased from Interchim (phalloidin488, FP-JO2830). β-casein from bovine milk (>98% pure, C6905), methylcellulose (M0512 4000 cPs) and other reagents were purchased from Sigma-Aldrich.

*Protein purification*. Muscle actin was purified from rabbit muscle and isolated in monomeric form in G-buffer (5 mM Tris-Cl\ pH 7.8, 0.1 mM CaCl2, 0.2 mM ATP, 1 mM DTT, 0.01% NaN3) as previously described^73^. Recombinant mouse IRSp53 I-BAR domain was purified as previously described^74^.

*GUV wetting on actin fibers*. All the steps were performed at room temperature. GUVs composed of DOPC supplemented with 0.5 mole% BODIPY TR ceramide and 0.5-1mole% DSPE-PEG(2000)-biotin were prepared by using polyvinyl alcohol (PVA) gel-assisted method in a sucrose buffer (200 mM) as described previously^75^. Muscle filamentous actin (F-actin) was pre-polymerized in an actin polymerization buffer (F-buffer, 5 mM Tris-Cl^-^, pH 7.8, 100 mM KCl, 0.2 mM EGTA, 1 mM MgCl2, 0.2 mM ATP, 10 mM DTT, 1 mM DABCO, 0.01% NaN3) in the presence of biotin-actin at a biotin-actin:actin molar ratio of 1:10, for at least 1hr. To visualize actin, F-actin was labeled with an equal molar amount of phalloidin488. Glass coverslips were bath sonicated with 2% Hellmanex for at least 30 min, rinsed with MilliQ water, sonicated with 1M KOH, and finally sonicated with MilliQ water for 20 min. Experimental chambers were assembled by placing a silicon open chamber on the cleaned coverslip. The glass substrate of the chamber was incubated with IRSp53 I-BAR domain proteins for at least 5 min, rinsed with F-buffer, incubated with β-casein (5 mg.mL^-1^) for at least 5 min, rinsed with F-buffer, incubated with F-actin at a concentration of 3 μM and in the presence of 0.3% methylcellulose for at least 15 min, rinsed with F-buffer, and finally incubated with NeutrAvidin (1 μM) for at least 10 min followed by rinsing with F-buffer. GUVs suspension in F-buffer were added into the chamber and incubated for at least 30 min before observation. To favour actin bundle formation, F-buffer was supplemented with around 20 mM MgCl2 when adding F-actin into the chamber. Samples were observed with an inverted spinning disk confocal microscope (Nikon eclipse Ti-E equipped with a 100X oil immersion objective and a CMOS camera, Prime 95B, Photometrics).

### Cryo-electron microscopy on vesicles with adhesive nanofibers

All steps were performed at room temperature. Vesicles were prepared by drying 0.3 mg of lipids dissolving in chloroform (DOPC/ 0.5 mole% DSPE-PEG(2000)-biotin) under nitrogen gas and then vacuum for at least 1 hr, followed by adding 300 μL of vesicle buffer (60 mM NaCl and 20 mM Tris pH 7.5) for resuspension, and finally a vortex mixing for 10 sec.

A 1.5 μM of F-actin containing 10 mole% biotin-actin in F-buffer was incubated with MgCl2 at a final concentration of 20 mM for 20 min to favor bundle formation, followed by incubating with NeutrAvidin at 0.7 μM for 5 min. 20 μL of vesicles were incubated with 20 μL of the pre-bundled F-actin for at least 1 hr before vitrification. The samples were vitrified on copper holey lacey grids (Ted Pella) using an automated device (EMGP, Leica) by blotting the excess sample on the opposite side from the droplet of sample for 4 seconds in a humid environment (90 % humidity). Imaging was performed on a LaB6 microscope operated at 200 kV (Technai G2, FEI) and equipped with a 4KX 4K CMOS camera (F416, TVIPS). Automated data collection for 2D imaging were carried out with the EMTools software suite.

### Cell culture on anodized aluminum oxide (AAO) membranes (or Anodiscs)

13 mm Whatman Anodiscs with 100 nm pores (GE Healthcare) were coated for 30 min with 0,1 μg.mL^-1^ APP3 in 10 mM HEPES pH 7.4, rinsed three times in 0,1M HEPES and once with distilled water, and then coated for 30 min with 100 μM BCN-RGD in PBS and rinsed three times in PBS ^33^. 1,5.10^5^ HUVEC were seeded per disc and cultivated overnight. The next day, cell culture medium was renewed with or without 100 nm cytochalasin D for 20 min at 37°C and cells were fixed in 2.5% glutaraldehyde (final concentration). Cells were then processed as for SEM.

### Preparation of native basal membranes from mouse mesentery

All experiments were conducted in accordance with the European Directive 2010/63/EU and national regulation for the protection of vertebrate animals used for experimental purposes (Decree 2013-118). Mice were kept in the Specific Pathogen Free (SPF) animal house of Institut Curie for breeding. Mesentery was isolated from a two-months old female C57BL/6 mouse according to previously described protocol^76,77^. Prior to mesentery isolation, the polyester membrane of a 6.4 mm diameter transwell insert (Corning) was removed using a scalpel. While holding the mouse intestine with tweezers, the mesentery was stretched and glued on the insert using surgical glue (3M Vetbond). Mesentery was washed with cold PBS supplemented with 1% Antibiotic-Antimycotic (Life Technologies), and incubated with 1M ammonium hydroxide (NH4OH, Sigma-Aldrich) for 40 min at RT to remove all cells present in the mesentery. Mesentery was washed three times with PBS supplemented with 1% Antibiotic-Antimycotic and stored at 4°C prior to seeding with HUVEC cells. Basal membrane with and without endothelial cells were processed for SEM as described above.

### Estimation of the binding energy for *Pseudomonas aeruginosa* T4P

We found an AFM study in which rupturing of type IV pili from *Pseudomonas aeruginosa* was performed^78^. In this article, there are two experiments where the rupture force was measured at two given velocities (see Figure 5 of the paper). Assuming a single barrier in the energy landscape, we extracted the binding energy *U* by using the following equation^79^

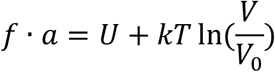

where *f* is the rupture force, *a* is the molecular length, *U* is the adhesion energy, *V* is the loading velocity and *V_0_* is a typical thermal velocity (in the order of 10 m.s^-1^). From the experimental data *f = 250 pN* for *V* = *1 μm. s^-1^* and *f =* 500 *pN* for *V* = *10 μm. s*^-1^, we found *U* ~ 18 *k_B_T*, which is consistent with the claiming of P.-G. de Gennes that for most practical separation experiments *U ~ 15 k_B_T*^79^.

### Statistical analysis

Data were analyzed in the Prism software (GraphPad). For comparison of exact values measured in multiple individual bacteria over several independent experiments, the data sets were first tested for normality with a D’ Agostino & Pearson test and then tested for statistical difference accordingly, either by an unpaired Mann-Whitney test (no normality) or an unpaired *t* test (normality). For comparison of mean values pooled over several independent experiments, statistical difference was tested with a paired *t* test. *, P<0.05; **, P<0.01.

### Data availability

The data that support the findings of this study are available from the corresponding author upon reasonable request

---

**Movie S1.**
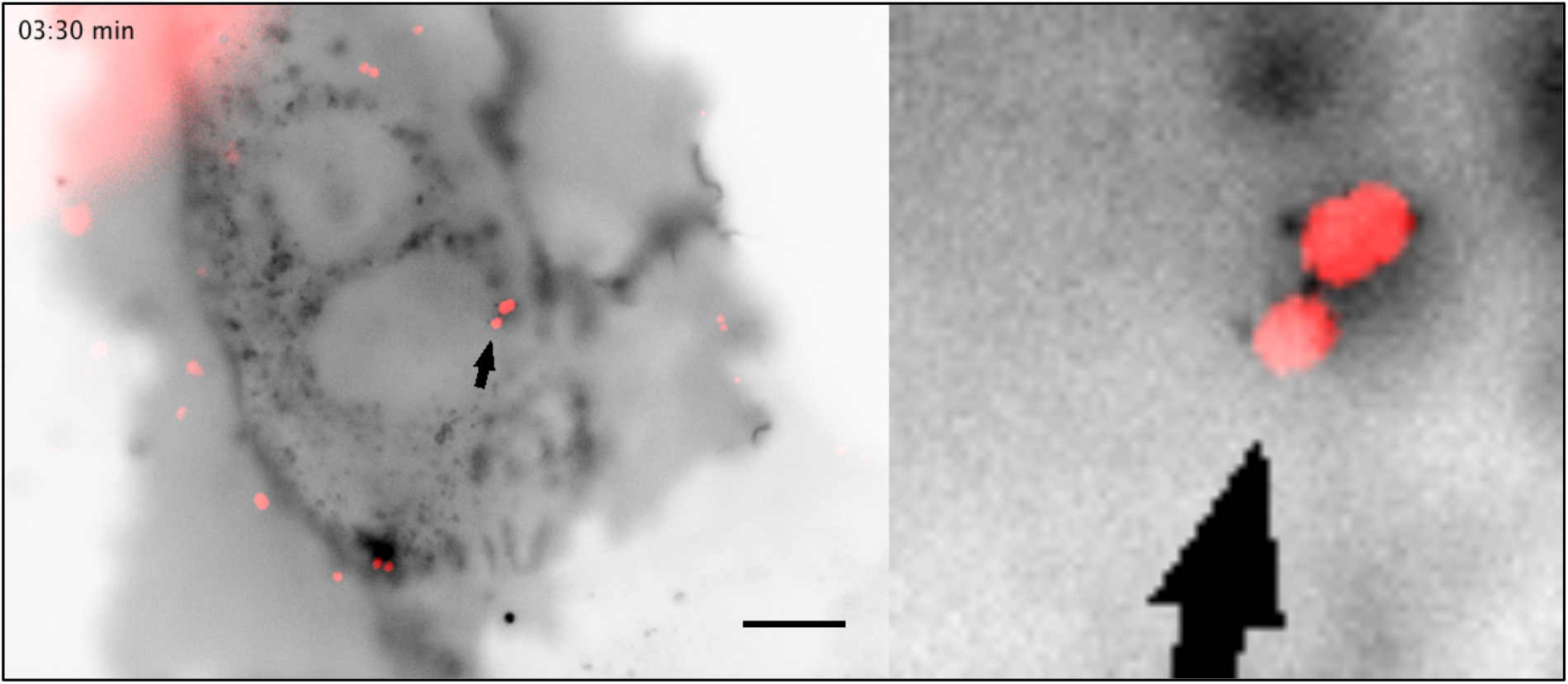
Oblique illumination live cell imaging of a HUVEC expressing the membrane marker GFP-F (inverted contrast) infected by Λ¾;-iRFP. A single focal plan is shown. Plasma membrane protrusions from the host cell are visible as discrete bright dots surrounding the bacterial bodies. Scale bar, 10 μm. Representative of several events in n>10 independent experiments.

**Movie S2.**
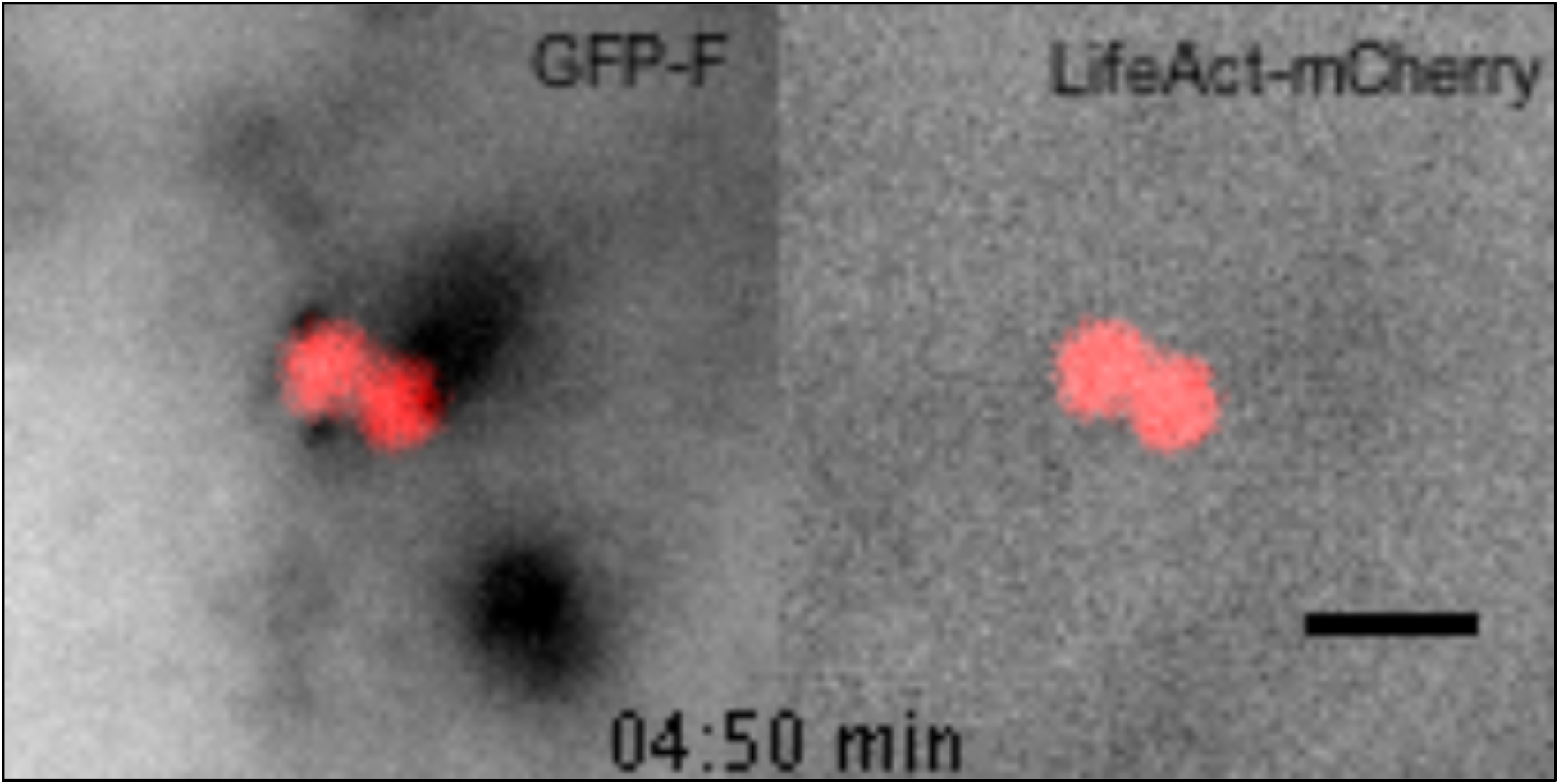
Oblique illumination live cell imaging of a HUVEC expressing the membrane marker GFP-F and LifeAct-mCherry infected by Λ¾?-iRFP. A single focal plan is shown. Plasma membrane protrusions from the host cell are visible as discrete bright dots surrounding the bacterial bodies. No accumulation of LifeAct-mCherry is observed. Scale bar, 2 μm. Representative of several events in n=2 independent experiments.

**Movie S3.**
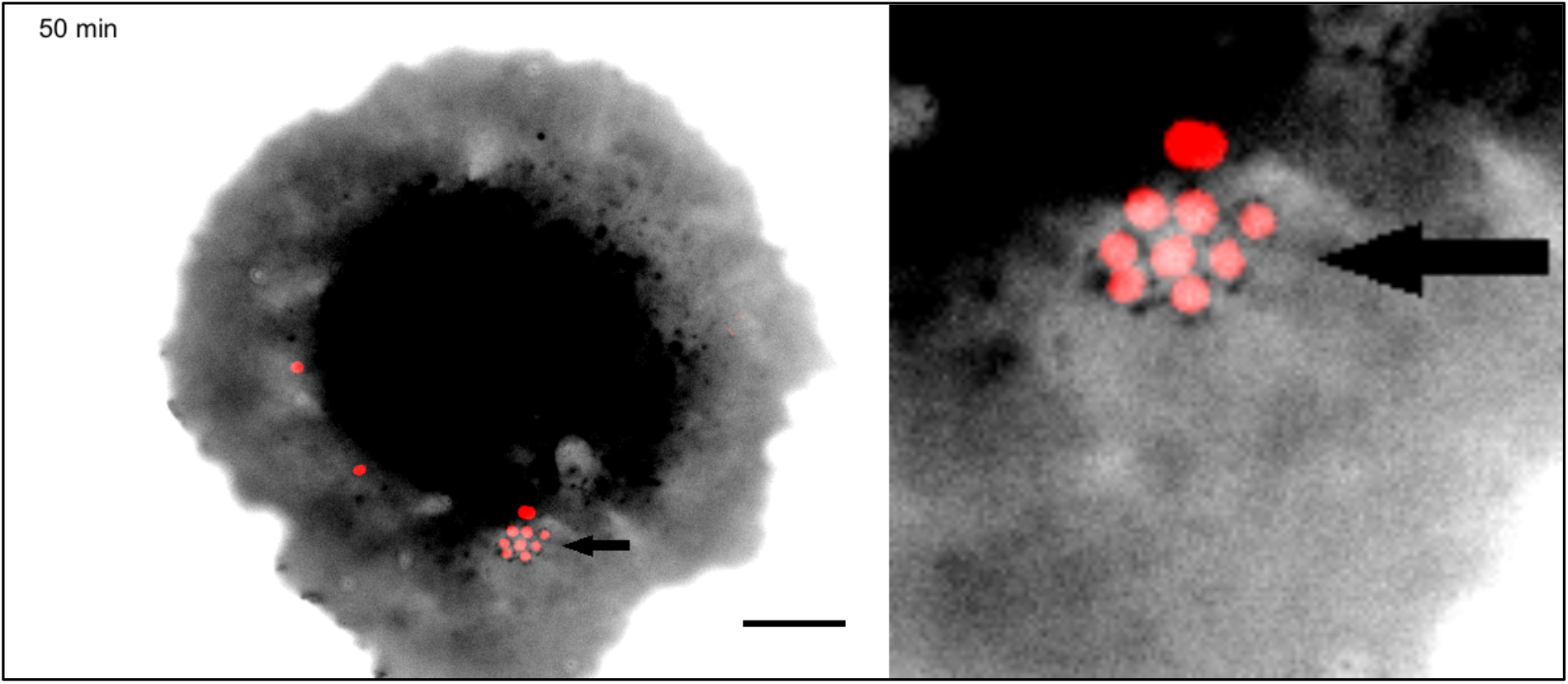
Oblique illumination live cell imaging of a micropatterned HUVEC expressing the membrane marker PM-EGFP (inverted contrast) and infected by Λ¾;-iRFP. A single focal plan is shown. Plasma membrane protrusions can be followed over two bacterial divisions. Scale bar, 10 μm. Representative of n=2 independent experiments.

**Movie S4.**
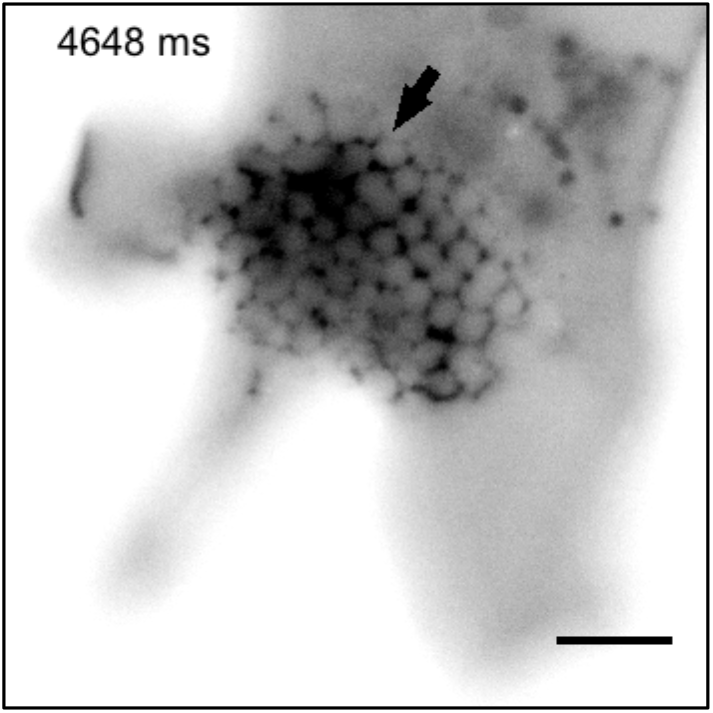
Oblique illumination live cell high speed imaging of a HUVEC expressing GFP-F (inverted contrast) infected with a pre-formed bacterial aggregate of Λ¾;-iRFP (not visible on the movie, please refer to Fig. 2). A single focal plan is shown. The plasma membrane protrusions barely move over 10 seconds, except at the aggregate periphery. Scale bar, 5 μm. Representative of several events in n>10 independent experiments.

**Movie S5.**
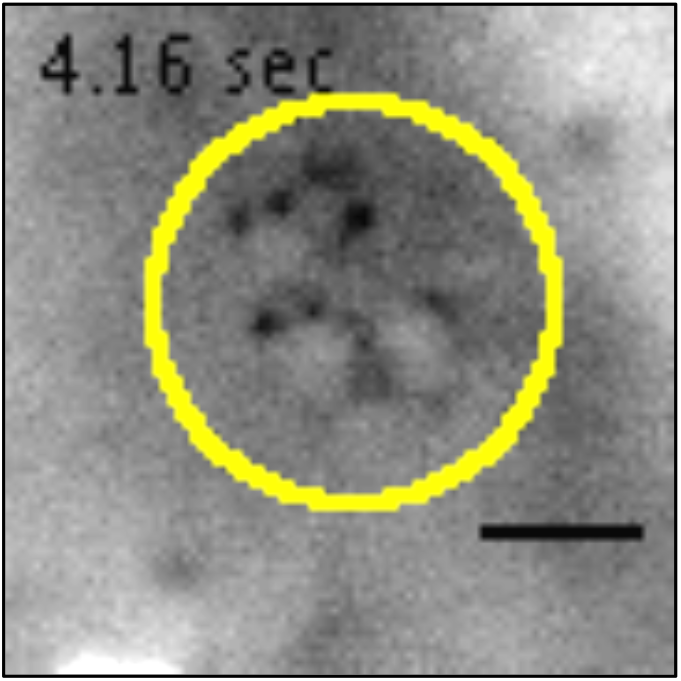
Oblique illumination live cell high speed imaging of a HUVEC expressing GFP-F (inverted contrast) infected with wild-type Λ¾;-iRFP bacteria. In this experimental setting, only one channel can be recorded at high speed. The position of 3 bacteria, denoted by a yellow ellipse, was assessed before recording of the GFP-F channel. Plasma membrane protrusions from the host cell are visible as discrete bright dots. Scale bar, 2 μm. Representative of several events in n=10 independent experiments.

**Movie S6.**
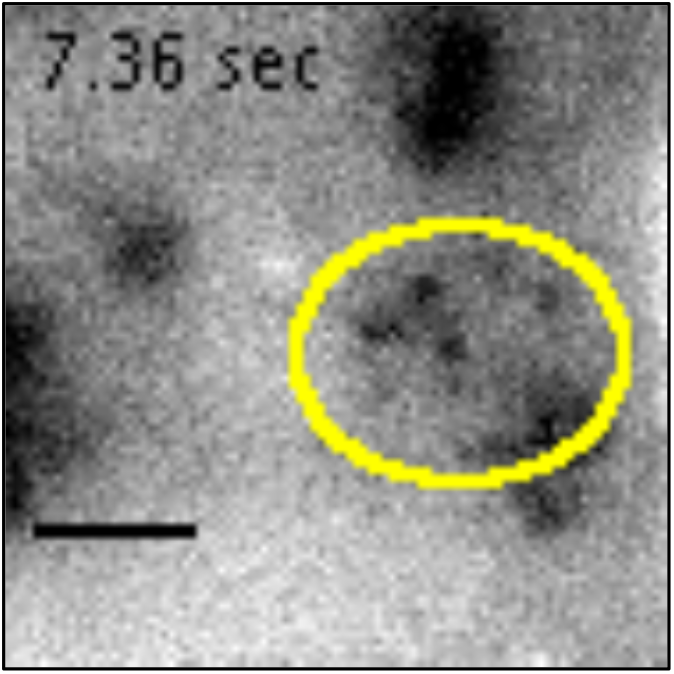
Oblique illumination live cell high speed imaging of a HUVEC expressing GFP-F and treated for 20 min with 100nM cytochalasin **D** (inverted contrast) infected with Λ¾;-iRFP bacteria. In this experimental setting, only one channel can be recorded at high speed. The position of 2 bacteria, denoted by a yellow ellipse, was assessed before recording of the GFP-F channel. Plasma membrane protrusions from the host cell are visible as discrete bright dots. Scale bar, 2 μm. Representative of several events in n=3 independent experiments.

**Movie S7.**
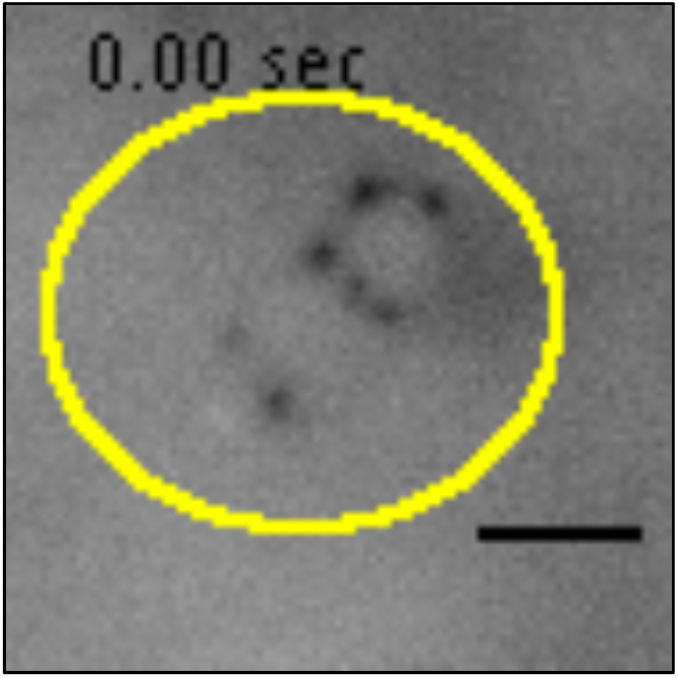
Oblique illumination live cell high speed imaging of a HUVEC expressing GFP-F (inverted contrast) infected with /;/V’ /-iRFP bacteria which are deficient for T4P retraction. In this experimental setting, only one channel can be recorded at high speed. The position of 2 bacteria, denoted by a yellow ellipse, was assessed before recording of the GFP-F channel. Plasma membrane protrusions from the host cell are visible as discrete bright dots. Scale bar, 2 μm. Representative of several events in n=3 independent experiments.

